# A prioritized and validated resource of mitochondrial proteins in *Plasmodium* identifies leads to unique biology

**DOI:** 10.1101/2021.01.22.427784

**Authors:** Selma L. van Esveld, Lisette Meerstein-Kessel, Cas Boshoven, Jochem F. Baaij, Konstantin Barylyuk, Jordy P. M. Coolen, Joeri van Strien, Ronald A.J. Duim, Bas E. Dutilh, Daniel R. Garza, Marijn Letterie, Nicholas I. Proellochs, Michelle N. de Ridder, Prashanna Balaji Venkatasubramanian, Laura E. de Vries, Ross F. Waller, Taco W.A. Kooij, Martijn A. Huynen

**Affiliations:** Center for Molecular and Biomolecular Informatics, Radboud Institute for Molecular Life Sciences, Radboudumc, Nijmegen, the Netherlands; Radboud Center for Mitochondrial Medicine, Radboudumc, Nijmegen, the Netherlands; Radboud Institute for Health Sciences, Radboudumc, Nijmegen, the Netherlands; Department of Medical Microbiology, Radboudumc Center for Infectious Diseases, Radboud Institute for Molecular Life Sciences, Radboudumc, Nijmegen, the Netherlands; Department of Biochemistry, University of Cambridge, Cambridge, UK; Theoretical Biology and Bioinformatics, Science for Life, Utrecht University, Utrecht, the Netherlands; Laboratory of Molecular Bacteriology (Rega Institute), Department of Microbiology, Immunology and Transplantation, KU Leuven, Leuven, Belgium

## Abstract

*Plasmodium* species have a single mitochondrion that is essential for their survival and has been successfully targeted by anti-malarial drugs. Most mitochondrial proteins are imported into this organelle and our picture of the *Plasmodium* mitochondrial proteome remains incomplete. Many data sources contain information about mitochondrial localization, including proteome and gene expression profiles, orthology to mitochondrial proteins from other species, co-evolutionary relationships, and amino acid sequences, each with different coverage and reliability. To obtain a comprehensive, prioritized list of *Plasmodium falciparum* mitochondrial proteins, we rigorously analyzed and integrated eight datasets using Bayesian statistics into a predictive score per protein for mitochondrial localization. At a corrected false discovery rate of 25%, we identified 445 proteins with a sensitivity of 87% and a specificity of 97%. They include proteins that have not been identified as mitochondrial in other eukaryotes but have characterized homologs in bacteria that are involved in metabolism or translation. Mitochondrial localization of seven *Plasmodium berghei* orthologs was confirmed by epitope labeling and co-localization with a mitochondrial marker protein. One of these belongs to a newly identified apicomplexan mitochondrial protein family that in *P. falciparum* has four members. With the experimentally validated mitochondrial proteins and the complete ranked *P. falciparum* proteome, which we have named PlasmoMitoCarta, we present a resource to study unique proteins of *Plasmodium* mitochondria.

## Introduction

Although all mitochondria evolved from a single endosymbiotic event (1), they display a large variety among the eukaryotes. For example, while in most species the mitochondrial organelle still retains its genome, the number of proteins encoded varies from 100 in *Andalucia godoyi* (2) to three in myzozoan species like *Plasmodium falciparum* that only encode cytochrome b, cytochrome c oxidase subunits 1 and 3, and two fragmented rRNAs (3). Moreover, mitochondria-like organelles of a variety of anaerobic species have lost the organellar genome altogether (4). Also mitochondrial proteomes vary broadly in size, ranging from e.g. ~1500 different proteins in mammalian mitochondria (5) to a few members of only a single biochemical pathway found in some mitosomes (6). Within this variety, the *P. falciparum* mitochondrion, based on the size of its genome, the size of the nuclear genome, and the complexity of its oxidative phosphorylation, is predicted to have a relatively small proteome. Although several mitochondrial pathways have already been identified in *P. falciparum* (7), and databases like PlasmoDB contain valuable information about mitochondrial localization of individual proteins (8), a complete resource that weighs and combines all the relevant information about possible mitochondrial localization is lacking.

*Plasmodium* mitochondria are particularly interesting because their function is variable between different life-cycle stages. In the sexual blood stage, functional oxidative phosphorylation is essential for colonization and development inside the mosquito hosts (9, 10), while in asexual blood stages the only essential function of the respiratory chain is to recycle ubiquinone for pyrimidine biosynthesis (11). This is further highlighted by the observation that in asexual blood-stage parasites no cristae are detectable in the mitochondrial membrane, while in gametocyte mitochondria cristae-like structures that typically accumulate respiratory chain complexes are observed (10, 12). A recent complexome profiling study demonstrated that the level of complex V components in asexual blood-stage parasites was only 3% of the level in gametocytes (12). Another intriguing aspect of mitochondria in *Plasmodium* and closely related species is that they have a rather unique FeS cluster assembly pathway (13). From this pathway the frataxin protein, an FeS assembly protein that occurs even in mitosomes (14) appears to be missing in *Plasmodium* (15).

Knowledge on the mitochondrial proteome could provide viable targets for the development of new drugs against this deadly parasite, as apicomplexan mitochondrial proteomes contain unique proteins that are not present in the animal hosts (16). Recently for example, an unusual prohibitin-like protein, PHBL, was identified in *Plasmodium berghei* that might provide a transmission-blocking drug target as it is unique to Myzozoa and *phbl*^−^ parasites fail to transmit (17). Furthermore, respiratory protein complex III has proven to be a good drug target (18) and inhibitors to the mitochondrial enzymes dihydroorotate dehydrogenase (DHODH) and NADH type II oxidoreductase (PfNDH2) have been identified (19).

For years, it has been difficult to perform proteomics experiments to identify the *Plasmodium* mitochondrial proteome due to the lack of organelle isolation methods that reliably separate mitochondria from the apicoplast (20). The definition of mitochondrial functions relied heavily upon homology studies (7, 21) and microscopic examination of individual proteins (Supplemental Table 1). With the use of two biotin tagging approaches, 422 mitochondrial matrix proteins were identified in *Toxoplasma gondii*, a species related to *Plasmodium* that also has an apicoplast and a mitochondrion (16). Besides these mitochondria-targeted approaches, hyperLOPIT, a whole-cell biochemical fractionation technique, defined the subcellular localization of the *T. gondii* proteome. This provided among others a set of 220 soluble and 168 membrane-bound *T. gondii* mitochondrial proteins (22). Alongside these proteomic data, there is a wealth of other data sources providing information on mitochondrial localization in *P. falciparum*. The challenge remains to use and combine these heterogeneous datasets to reliably predict the mitochondrial proteome.

Here, we integrated eight datasets containing proteomic, gene expression, orthology to mitochondrial proteins from other species, phylogenetic distribution, and amino acid sequence data in a Bayesian manner as has been applied before such as for the definition of the mitochondrial proteome of humans (13) and of developmental stage specific proteins in Plasmodium(23, 24). We used a naïve Bayesian classifier to combine the data. This method exploits the relative strengths of the various datasets while maintaining transparency about the contribution of each dataset to the final Bayesian posterior probability score for being mitochondrial. For each protein, the contribution of each dataset to the prediction was determined using gold standards of *Plasmodium* proteins that are either known to be mitochondrial or non-mitochondrial. After data integration, we obtained a list of 445 P. falciparum mitochondrial proteins. We experimentally validated these predictions with seven proteins of varying probability scores, and collectively this predicted proteome indicates that the *P. falciparum* mitochondria are unique relative to mitochondria of model organisms.

## Materials and Methods

### Animal experiments

All animal experiments were performed in accordance with the Dutch Experiments on Animals Act (Wod), the Directive 2010/63/EU from the European Union and the European ETS 123 convention and approved by the Radboud University Animal Welfare Body (IvD) and Animal Experiment Committee (RUDEC; 2015-0142), and the Central Authority for Scientific Procedures on Animals (CCD; AVD103002016424). In this study, we used outbred male and female NMRI mice (Envigo).

### Nuclear encoded reference proteome

All datasets used in this study were mapped to the *P. falciparum* 3D7 reference proteome version 3.1 from the Sanger Institute, downloaded December 2017. This version, including isoforms, contains 5431 proteins encoded by 5357 genes. As we are interested in the proteins encoded by nuclear genes, all apicoplast and mitochondrial encoded proteins were excluded, leaving a reference of 5324 genes. When a dataset contains informational data points for two or more protein isoforms of one gene, the value chosen for that gene was the one that comes closest to the expected values for mitochondrial proteins in that specific dataset.

### Assembly of positive and negative lists for benchmarking

Bayesian data integration depends on gold standards of proteins known to be mitochondrial or non-mitochondrial. We constructed two gold standards to assess the predictive value of the individual datasets. Furthermore, we built two alternative sets for the construction of the CLIME (co-evolution) and WICCA (co-expression) datasets to prevent that the standards chosen to train those two predictors were also used to evaluate them. Thus we avoid circular arguments in the data integration. These four sets are unique, have no overlapping genes, and are available in Supplemental Table 1. To construct these lists, we commenced with a systematic review of all literature available via PubMed (as per 30-08-2018) using the following broad search string “(mitochondri* OR apicoplast OR plastid) AND (plasmodium OR malaria)”. Given the intimate functional and physical relation between mitochondrion and apicoplast, we reasoned that the positive gold standard list should only include genes encoding proteins that had been unambiguously and singularly assigned to the mitochondrion through fluorescence microscopy and co-localization with confirmed mitochondrial markers or via immune-electron microscopy (54 genes). Any proteins for which multiple studies showed different results were excluded. To facilitate discrimination from the apicoplast, we also made a gold standard apicoplast list (72 genes). Finally, all proteins that were unambiguously non-mitochondrial as demonstrated in the same papers (170 genes) were combined with the apicoplast-positive list to form the basis for the negative gold standard. This negative set was then complemented with an additional 346 non-overlapping genes encoding non-apicoplast and non-mitochondrial proteins identified through extensive literature review and published online by the Ralph lab (25). The 588 genes encoding non-mitochondrial proteins were randomly assigned to the negative gold standard (294 genes) that was used for the Bayesian data integration and the alternative negative gold standard (294 genes) that was used to train the CLIME and WICCA approaches. For the latter purpose, an alternative positive standard was compiled of 146 genes, non-overlapping with the first gold standard, but associated with one or more mitochondrial GO-terms (GO:0000275, GO:0005739-43, GO:0005746, GO:0005750, GO:0005753, GO:0005758, GO:0005759, GO:0006122, GO:0006839, GO:0006850, GO:0031966, GO:0033108, GO:0042775, GO:0044429, GO:0044455).

### Datasets to predict mitochondrial and non-mitochondrial proteins

To predict mitochondrial proteins and non-mitochondrial proteins, we collected and constructed eight datasets (Table 1) that contain features typical of either mitochondrial or apicoplast proteins. Some proteins, like lipoate protein ligase 2 (PF3D7_0923600) (26) and serine hydroxymethyltransferase isoforms (PF3D7_1235600) (27), are dual localized to the mitochondrion and apicoplast, highlighting the need for the second category of datasets. Each dataset is described in detail below, including how the dataset was processed to obtain per *P. falciparum* protein a single score relevant to whether the protein is mitochondrial or not.

The size of datasets indicates the number of *Plasmodium* genes that could potentially have been measured in this dataset. Fractions indicate the actually identified genes of this set that fall in bins with a positive mitochondrial score (Fig 1A). P values (Fisher’s exact test) indicate overrepresentation of identified gold standard genes compared to random expectation.Notice that in the “non-Apicoplast proteins” and in the “Absence of a Cryptosporidium ortholog” these P values reflect the over representation of mitochondrial proteins among the large sets of proteins to which these features apply.

**Table 1:** Input datasets for Bayesian integration.

| Dataset | Abbreviations used in figures | Size of dataset | Fraction identified (%) | Fraction gold identified (%) | P - value |
| --- | --- | --- | --- | --- | --- |
| <u>Dataset based on 'omics data</u> |  |  |  |  |  |
| Co-expression, built using in-house developed WICCA tool (rank $\leq 2400$) (https://wicca.cmbi.umcn.nl/) | Co-expression | 5310 | 45.6 | 83.0 | 2.0E-08 |
| <u>Non- Apicoplast proteins (28)</u> | BioID, gene expression | 5324 | 93.6 | 94.4 | 5.4E-01 |
| <u>Datasets based on ortholog evidence</u> |  |  |  |  |  |
| hyperLOPIT (TAGM.MCMC.joint.mitochondrion_max $\geq 4.78\text{E-}05$) (22) | hyperLOPIT | 2246 | 15 | 95.2 | 9.3E-32 |
| Mitochondrial ortholog | Mito ortholog | 5280 | 8.7 | 75.5 | 2.6E-32 |
| Evolutionary inference, built using CLIME (NN score $\leq -1.5$) (29) | CLIME | 4687 | 9.0 | 38.5 | 6.9E-09 |
| Absence of a *Cryptosporidium* ortholog (score $\leq 4$) | Crypto ortholog | 5280 | 67.5 | 77.4 | 7.9E-02 |
| <u>Datasets based on amino acid sequence evidence</u> |  |  |  |  |  |
| Mitochondrial targeting signal, built using PFMpred tool (SVM $\geq -0.67$) (30) | Mitochondrial TS | 5324 | 59.5 | 83.3 | 1.4E-04 |
| Isoelectric point (Patrickios pl $\geq 4.2$)(28) | pl | 5324 | 55.1 | 83.3 | 1.1E-05 |

**Fig 1.**
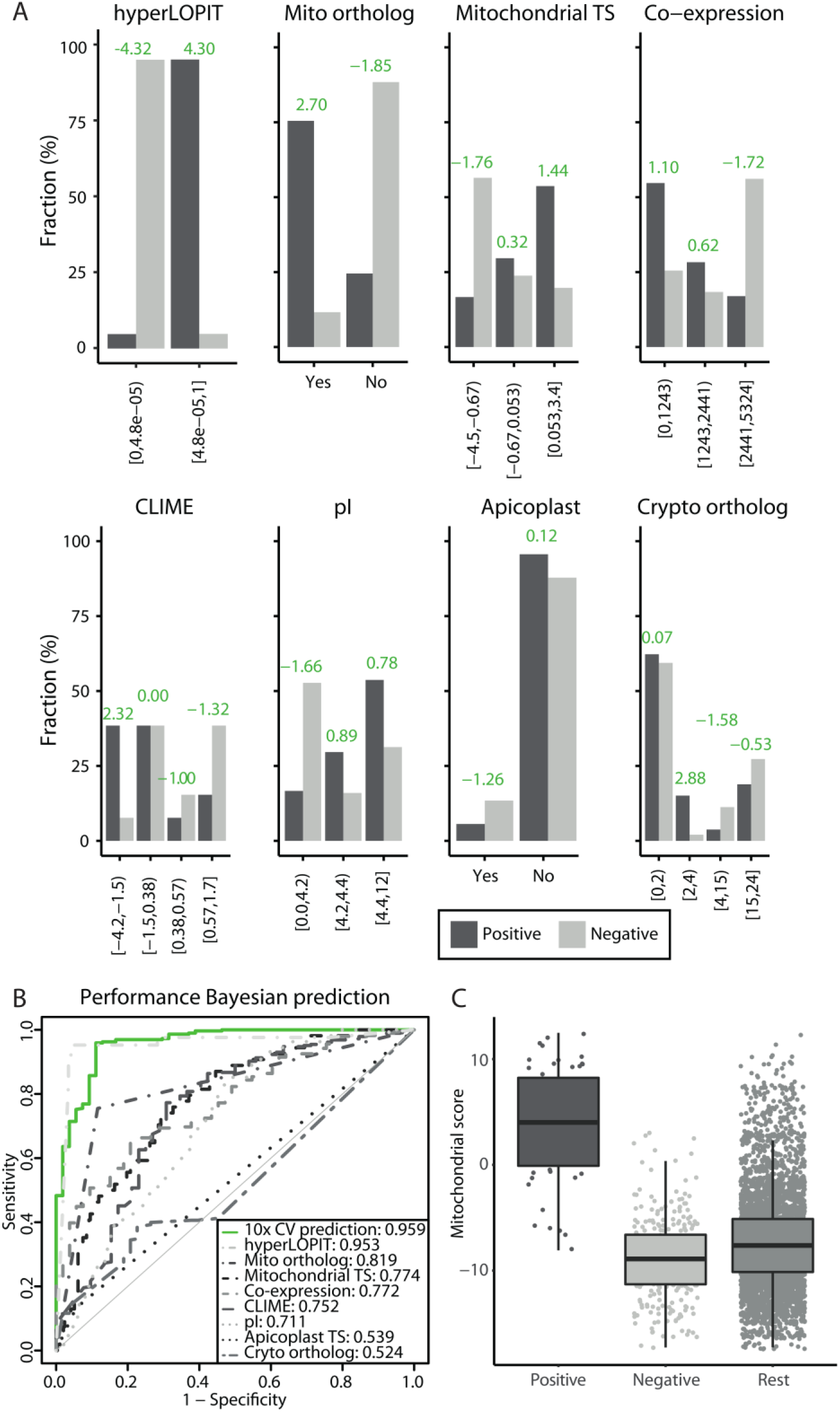
Predictive values of the individual datasets and the integrated ranked list. (A) Barplot of each dataset with the fraction of the gold standards on the y-axis and the bins on the x-axis. In green, the mitochondrial score per bin is indicated to show the predictive value of each bin. (B) ROC-curves comparing the performance to identify mitochondrial proteins of the prediction (10-fold cross-validated) to the performance of the individual datasets. For comparison, the values for the area under each curve are indicated in the legend. (C) Boxplot that visualizes the distribution of the gold-standard positives, negatives, and remainder of the proteome (x-axis) over the calculated mitochondrial score of the prediction (y-axis).

#### Dataset based on ‘omics data

### Co-expression

Co-expressed genes tend to code for proteins that functionally interact. For mammals, co-expression analyses have been successfully used in the discovery of mitochondrial proteins (5, 31). To determine if the expression pattern of a *Plasmodium* gene correlates with mitochondrial protein coding genes, we developed a co-expression tool: WeIghted Co-expression Calculation tool for plAsmodium genes (WICCA, available at https://wicca.cmbi.umcn.nl/). In short, the tool uses the co-expression of a group of input genes with each other to weigh 83 microarray experiments for their predictive value for that group. Datasets that show high co-expression of the input genes are more likely to be relevant for the system that the input genes are part of, and receive a higher weight. It then uses these weights to combine the expression datasets and calculate one co-expression score per gene with the input system, in this case, one score per gene for the level of co-expression with mitochondrial protein coding genes. We used the co-expression ranking obtained with the alternative positive standard of 146 genes as input for WICCA to weigh the co-expression datasets. The methods of this tool were based on the WeGET method (http://weget.cmbi.umcn.nl/, (32)), methodological details on WICCA can be found in the supplement.

### Bio-ID

In Bayesian data integration, “negative data sets” that e.g. predict that a protein is non-mitochondrial can be as valuable as positive ones. We used the set of 346 apicoplast proteins that are based on BioID data and gene expression data analyzed with a Neural Network (28) to reduce apicoplast protein contamination among the top scoring proteins.

#### Datasets based on ortholog evidence

### hyperLOPIT

Hyperplexed Localization of Organelle Proteins by Isotope Tagging (hyperLOPIT) is a proteomics technique that analyzes protein distributions upon biochemical fractionation. It enables the identification of the subcellular localization of thousands of proteins (33, 34). Barylyuk *et al.* used this technique on the apicomplexan *T. gondii* and, among others, classified proteins identified in all three hyperLOPIT experiments to be part of the mitochondrial soluble matrix and mitochondrial membrane. This classification was based on t-augmented Gaussian mixture models (TAGM) in combination with maximum a posteriori prediction (TAGM-MAP) and Markov-chain Monte-Carlo (TAGM-MCMC) methods (22). Marker proteins used for the classification were arbitrarily set to 1 in the class that they belong to and to 0 for all other classes. Using BLAST (35), or if that produced no homologs HHpred (36), in combination with selecting for best bidirectional hits, we determined a list of orthologs between *T. gondii* and *P. falciparum*. The 3,832 *T. gondii* proteins that got a TAGM-MCMC probability for mitochondrial soluble matrix and mitochondrial membrane were mapped to the corresponding *P. falciparum* ortholog when available. This resulted in a list of 2246 *P. falciparum* identifiers with two TAGM-MCMC probabilities, one for mitochondrial matrix localization and one for mitochondrial membrane localization. As the input datasets for the Bayesian integration should be independent, both TAGM-MCMC probabilities cannot be included as separate datasets. Therefore the maximum TAGM-MCMC probability (so either the soluble or the membrane value) for each protein was used to create one input hyperLOPIT dataset.

### Mitochondrial ortholog

We also used orthologs from more distantly related species than *T. gondii*, as also at larger phylogenetic distances subcellular localizations tend to be conserved (37). Using BLAST (35) (all versus all blast for proteomes with e-value of 100) in combination with OrthoMCL (38) (percent match cut of 50, e-value exponent cut off −5) to detect orthologs, or if that produced no result, HHpred (36) (default settings, e-value cut off 0.01, three iterations, only best-bi-directional hits included), one-to-one orthologs between *P. falciparum* and either *Homo sapiens*, *Arabidopsis thaliana* or *Saccharomyces cerevisiae* were determined. When the orthologous protein of at least one species was annotated to be mitochondrial in a published mitochondrial compendium (*H. sapiens* MitoCarta2.0 (5), *A. thaliana* (39), *S. cerevisiae* (40)), then that *P. falciparum* protein was included in the mitochondrial ortholog dataset.

### Evolutionary inference

Clustered by inferred models of evolution (CLIME) uses homology across model species (138 eukaryotic species and 1 prokaryotic outgroup) to identify evolutionary conserved clusters. We downloaded the pre-computed CLIME analyses of *P. falciparum* genes from http://gene-clime.org/ (29) and mapped the IDs to our reference proteome. We used the CLIME matrix listing the presence/absence of orthologs of *P. falciparum* in other species, with the alternative positive and negative sets as training data, to train a perceptron with two hidden layers to obtain a single score per protein (details in supplement). The resulting output scores on our test set formed the CLIME input for the Bayesian integration.

### Absence of a *Cryptosporidium* ortholog

Like *Plasmodium*, the genus *Cryptosporidium* belongs to the phylum of Apicomplexa. *Cryptosporidium* species lack an apicoplast and contain only a remnant mitochondrial-like organelle (41). Therefore, it can be expected that apicoplast proteins and many mitochondrial proteins will not have an ortholog in *Cryptosporidium*. MetaPhOrs is an online tool that contains orthology and paralogy predictions obtained from multiple phylogenetic trees (42) and was used to assess orthology for *P. falciparum* genes in three *Cryptosporidium* species (*hominis, parvum*, and *muris*). We calculated a combined score by multiplying two MetaPhOrs metrics, one for the confidence level (see below) and one for the number of hits between the four species, and used this combined score as input for the Bayesian integration. In detail, the MetaPhOrs results include a consistency score (ranging from 0 with no overlap to 1 if all trees contain the same protein relationship/orthology information) and an evidence level for the number of independent databases (the theoretical maximum is 13, but for our species combination it was four). The multiplication of consistency and evidence level results in an arbitrary score (range 0-4) that indicates the confidence in calling two proteins orthologs. *P. falciparum* proteins with a high score are more likely to have an ortholog in *Cryptosporidium*. Notice that there is some overlap with the Evolutionary inference as that includes one *Cryptosporidium* species. Nevertheless, *Cryptosporidium* species show quite some variation in the complexity of their mitosomes and the overlap in the predictions is limited (see below).

#### Datasets based on amino acid sequence evidence

### Mitochondrial targeting signal

The canonical mitochondrial import system requires an amphipathic α-helical N-terminal targeting sequence. As *Plasmodium* has a distinct amino acid usage pattern (43), a *P. falciparum* specific tool, PFMpred (30), was chosen to predict mitochondrial localization based on the amino acid sequence. This is a support vector machine (SVM) based tool and was used in split-amino-acid-composition mode (Matthews correlation coefficient 0.73) that allows separate calculations for the N-terminus, C-terminus, and remainder of the protein. The tool reports one SVM-score per protein and indicates whether the protein is predicted to be mitochondrial or not. The SVM-scores per gene are used as the mitochondrial targeting signal dataset.

### Isoelectric point

Nuclear encoded mitochondrial proteins need to cross the negatively charged mitochondrial membrane and need to function properly inside the mitochondrial environment that has a slightly higher pH compared to the cytosol (44). A density plot (Supplemental Fig 1) by Patrikios isoelectric points (45) of the human mitochondrial proteome (MitoCarta2.0 (5)) showed that mitochondrial proteins on average have a higher pI than the remainder of the proteome. Isoelectric points for the *P. falciparum* proteome were calculated using the Patrickios algorithm (46) and directly used as pI scores.

### Bayesian integration to predict mitochondrial proteins

Using the eight datasets described above and the gold standard evaluation sets, a mitochondrial score for each *P. falciparum* reference protein was calculated. This score is the logarithm of the odds that a protein is localized in the mitochondrion relative to the protein being localized somewhere else in the cell. For categorical data (mitochondrial ortholog, apicoplast targeting signal), the dataset is separated into bins that represent the two categories. Continuous data (the other six datasets) were binned in a systematic way using a custom python script (https://github.com/JordyCoolen/Binning) (settings: --bins 0 --score 1). In short, this script optimizes for each dataset the distribution of the bins such that the sum of log ratio scores of all bins combined (see below) is close to zero, as this will result in the best separation of positive and negative bins. The number of bins was varied to a maximum of five bins to achieve the most optimal separation without expanding the number such that there are too few proteins per bin to obtain reliable estimates of the log odds of that bin.

The fractions of gold standard proteins per bin are determined. The log ratio of these fractions determines the score for all other proteins in that respective bin. The mitochondrial score is based on the sum of log ratios of the individual datasets and is calculated as follows:

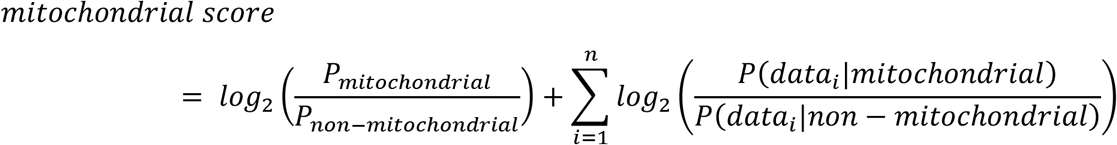

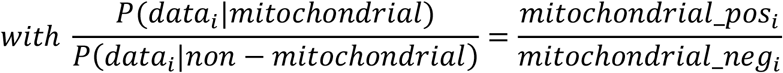

where mitochondrial_pos and mitochondrial_neg are the fractions of the positive and negative gold standard genes in sample i respectively. If there were no gold standard genes found in a certain bin, that positive or negative gold standard fraction was set to 0.5/’total number of negative set genes in the complete dataset’ to prevent division by zero and allow calculation of the log ratio. The O_prior_, log_2_(P_mitochondrial_/P_non-mitochondrial_), is based on the estimation that 536 proteins out of the 5357 protein coding genes (~10%) encode a mitochondrial protein. This is a conservative estimate when compared to single cell species like *Saccharomyces cerevisiae* with 16% mitochondrial proteins (47), or the more closely related *T. gondii* where 15% of proteins that could confidently be mapped are mitochondrial (48), while compared to all proteins that were mapped in that dataset the percentage of mitochondrial proteins is 10%. Note that the O_prior_ only affects the score per protein, it does not affect the relative ranking of potential mitochondrial proteins.

To assess the performance of the integration, a false discovery rate (FDR) was calculated. As this FDR depends on gold standard genes and as the ratio gold positives/negatives is not similar to the ratio of mitochondrial protein coding genes versus non-mitochondrial coding genes in the genome, the FDR was corrected (cFDR) using the following formula:

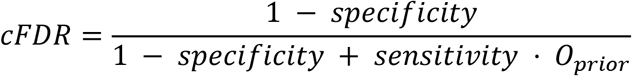

### Cross-validation

To assess the ability of the integrated predictor to discriminate known mitochondrial proteins from non-mitochondrial proteins, ten-fold cross-validation was performed. The gold standards (both negative and positive) were subsampled ten times, thereby creating ten sets of nine-tenth of the gold standard genes. Each gold standard gene was left out once in one of the ten sets. Data integration was performed with each of these ten sets and the ranks of the one-tenth left out gold standard genes were retrieved. The ten-fold cross-validated ROC curve was constructed based on those ranks.

### Intraspecies homologs

The human mitochondrial proteome has expanded since the divergence from yeast mainly due to gene duplications that create mitochondrial paralogs (37). It is therefore interesting to see if the predicted *P. falciparum* mitochondrial proteins have intraspecies homologs, and whether we can identify mitochondrial gene families. With HHpred (36), using default settings and a 1E-5 E-value cut-off, the homologs within the genome of *P. falciparum* were determined and included as a column in Supplemental Table 2.

### Validation of candidate proteins

To validate the predictions, we selected 14 genes with unusual or novel mitochondrial functions within the top 295 genes that fell within the 25% cFDR cut-off in the initial data integration. Note that during the project, the number of predicted mitochondrial proteins increased to 445 by including better orthology prediction and a better data set of likely apicoplast proteins. Of these 14 genes, 13 were selected based on maximum transcript levels during asexual blood-stage development of *P. berghei* strain ANKA cl15cy1 that exceeded 100 FPKM (49). In addition, we selected one particularly interesting gene, PF3D7_1004900 that had a lower transcript level. We used experimental genetics (50) to generate *P. berghei* lines expressing 3xHA-tagged copies of the selected proteins and colocalized them with an established mitochondrial marker (17) to determine their subcellular localization by immunofluorescence microscopy. Details on plasmid construction (successful for 13 out of 14 targets, excluding PF3D7_1142800) and the generation of the lines can be found as supplemental information. To assess colocalization, the freshly harvested transgenic parasites were allowed to settle on a poly-L-lysine coated coverslip for 10 minutes and then fixed with 4% EM-grade paraformaldehyde (Fisher Scientific) and 0.0075% EM-grade glutaraldehyde (Sigma-Aldrich) in microtubule stabilizing buffer (MTSB, 10 mM MES, 150 mM NaCl, 5 mM EGTA, 5 mM glucose, 5 mM MgCl2 pH 6.9) for 20 minutes (51). Next, parasites were permeabilized with 0.1% Triton X-100 in PBS for 10 minutes. Samples in which we imaged mOrange and mCherry were hereafter stained with DAPI (1:300) for 15 minutes and mounted on a microscope slide with VectaShield (Vector laboratories). All other samples were blocked in 3% FCS-PBS for 1 hour. Samples were incubated overnight at 4°C with rat anti-HA (1:500, ROAHAHA Roche) and chicken anti-GFP (1:1000, Thermo Fisher) antibodies and for 1 hour with 1:500 goat anti-rat Alexa fluor™ 594 (Invitrogen) and goat anti-chicken Alexa fluor™ 488 (Thermo Fisher) conjugated antibodies at room temperature. Nuclei were stained with DAPI (1:300) for 15 minutes and coverslips were mounted on a microscope slide with VectaShield (Vector laboratories). Samples were imaged with an SP8 confocal microscope (Leica, 63x oil lens) or LSM900 confocal microscope (Zeiss, 63x oil lens). Images were processed minimally and similarly with Fiji (52).

### Tools for data analysis

Plots, statistics, and calculations were performed with the R statistical package (53) and additional packages gplots (54), ggplot2 (55), ROCR (56), scales (57) and reshape (58). The separation of the datasets into bins was performed using a python script (see above).

## Results

We integrated eight features of proteomic, gene expression, orthology and amino acid composition data, to predict mitochondrial localization in *P. falciparum*. All the individual features had a predictive value for mitochondrial localization (Fig 1A and 1B), including, as expected, a negative score for the “*Cryptosporidium* ortholog” and the “apicoplast” datasets (Supplemental Fig 2A). Notice that for the co-expression feature it is the bin with the low-rank values that contains most mitochondrial proteins because this bin contains the proteins with the highest co-expression values. The best performing input datasets are the ones that translate mitochondrial localization between species: the hyperLOPIT dataset and the mitochondrial ortholog dataset, followed by the mitochondrial targeting signal and co-expression datasets (Fig 1B). Correlation analyses showed that the features are largely independent off each other, allowing us to use naïve Bayesian data integration (Supplemental Fig 2B). The highest correlations were observed between features that examine the phylogenetic distribution of the proteins: CLIME and *Cryptosporidium* ortholog datasets. By integrating all data, we achieved both a better sensitivity and specificity in predicting mitochondrial proteins with an AUC of 0.959 than we achieved for individual data sets (Fig 1B). The performance of the hyperLOPIT dataset on its own approximates the integrated prediction with an AUC of 0.953 (Fig 1B). However, this dataset contains information on only 2246 (42%) of the nuclear encoded *Plasmodium* proteins, highlighting the value of additional datasets and their integration to predict a complete mitochondrial proteome. The distribution of the log-odds ratios (Fig 1C) showed that the mitochondrial score separated the positive and negative gold standards well from each other and that there are additional proteins with a high score that were not part of either, which are potential new mitochondrial proteins. We use a cFDR of 25% (Fig 2A) as a threshold for highly probable new mitochondrial proteins and identify 445 proteins (Fig 2B) with a sensitivity of 87% and a specificity of 97%. The ranked list of the complete nuclear encoded proteome is available in Supplemental Table 2.

**Fig 2.**
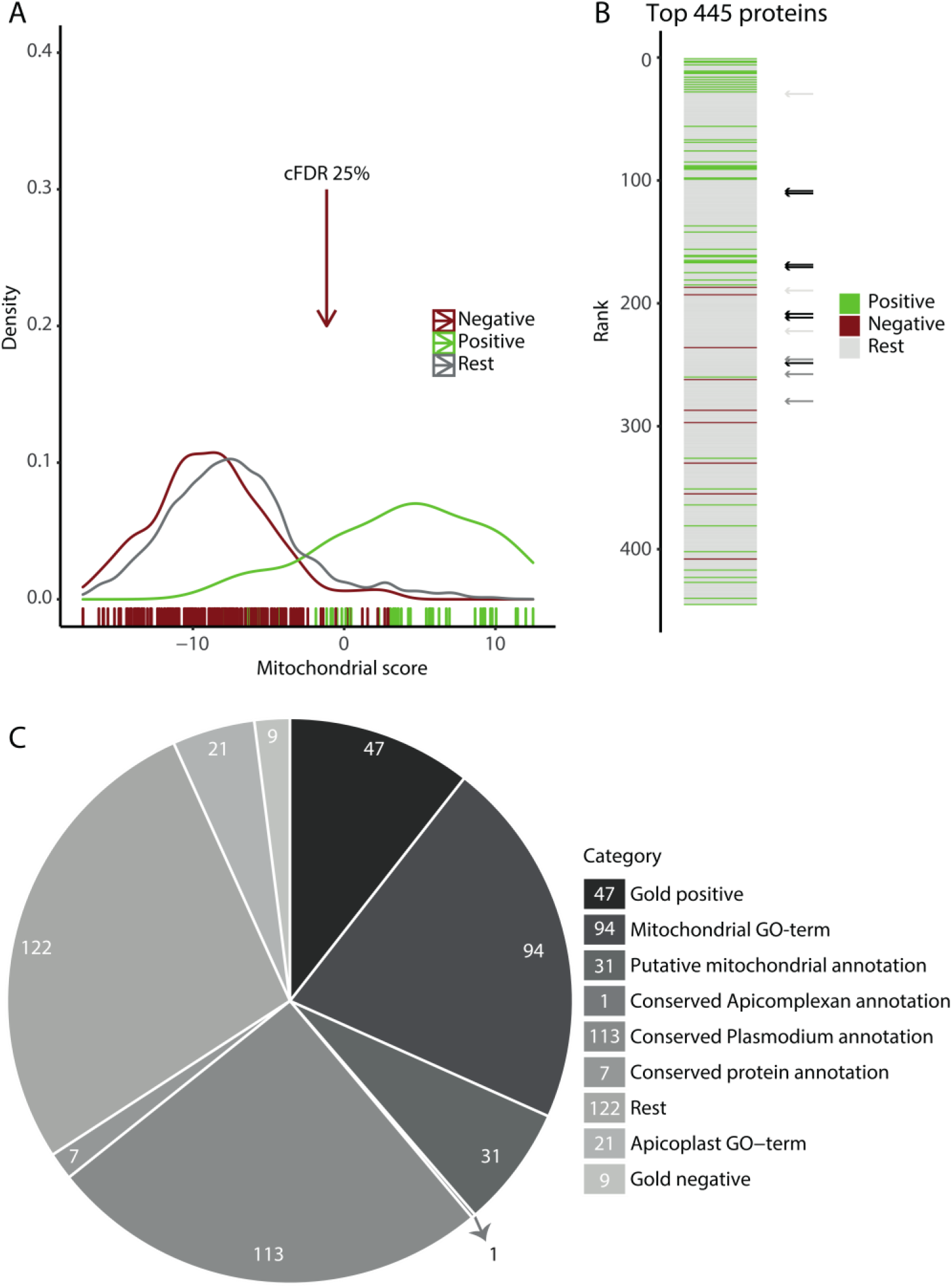
Integration produces a ranked proteome with 445 likely mitochondrial proteins at a 25% cFDR-cut. (A) Density plot of the mitochondrial score with a colored bar at the bottom indicating the scores of individual gold standards. The arrow indicates the 25% cFDR cut-off. (B) The top 445 proteins falling in the 25% cFDR cut-off ranked on their mitochondrial score with color indicating if the protein is part of a gold standard. Black arrows indicated the ranking of the seven candidate proteins that we experimentally confirmed to be mitochondrial (Fig 3A and 3B), dark grey arrows indicate the ranking of the three proteins with suggested mitochondrial localization (Fig 3C), and light grey arrows the ranking of the three proteins with inconclusive results (Supplemental Fig 4B). (C) Categorical representation of the 445 proteins identified at the 25% cFDR cut-off. First, genes were separated into categories based on the gold standard, the remainder on mitochondrial and apicoplast GO term, and subsequently the remainder on gene function annotations.

### Comparison of the Toxoplasma and Plasmodium mitochondrial proteomes

Overall, the hyperLOPIT dataset contains 388 *T. gondii* mitochondrial proteins with a TAGM-MCMC location probability of 99% or higher. Of those, 296 have a *P. falciparum* ortholog and of these orthologs, we find 273 in our predicted mitochondrial proteome. Most proteins that we predicted to be mitochondrial that were not supported by the *T. gondii* data are proteins that do have orthologs in *T. gondii*, for which there were however no hyperLOPIT data (95 proteins). There are furthermore 38 proteins in the set of 445 that do not have *T. gondii* orthologs. It should be noted that these do not include the six *P. falciparum* mitochondrial ribosomal proteins that, based on Blast searches, were deemed to be absent from *T. gondii* (59), as for those we could identify orthologs using HHpred (Table S2), underlining the relevance of sensitive homology detection. Finally, there are 39 proteins of which orthologs are present in the hyperLOPIT dataset but fell outside the 99% cut off for mitochondrial localization that was used in that study, 26 of these did still have a mitochondrial localization as the most probable location and 13 had a different predicted location in *T. gondii* (flagged in Supplemental Table 2). Manual examination of those 13 inconsistencies reveals proteins like PF3D7_1125300/mitochondrial RNA polymerase that in the hyperLOPIT data has been observed in the nucleolus, but also PF3D7_1446400/pdhB that was predicted to be mitochondrial based on orthology with pdhB in e.g. *H.sapiens* but is localized in the apicoplast in *Plasmodium(60)*.

#### Validation of candidates

We aimed to validate the mitochondrial candidate list by tagging orthologs of the selected unusual or unique predicted mitochondrial proteins in the efficient transfection model *P. berghei* (see the section ‘new mitochondrial proteins’ for the *P. falciparum* orthologs of these proteins). Mitochondrial localization was assessed by fluorescent microscopy using an experimental genetic approach developed in our lab (17). This method allows the endogenous tagging of a target protein with a combined fluorescent and epitope tag, while simultaneously introducing a strong mitochondrial targeted GFP marker by fusion of the promoter and N terminus of HSP70-3 (PBANKA_0914400) to GFP. Initially, we tagged 6 proteins with a mOrange-3xHA tag. Live imaging revealed very poor and undefined signals (data not shown). Next, we fixed blood samples and stained them with anti-HA antibodies. Using this approach, only PBANKA_0310100 and PBANKA_1203200 were convincingly shown to localize to the mitochondrion (Fig 3A), while other proteins demonstrated mostly undetectable or undefined signals (Supplemental Fig 4A). Since many of the selected proteins are relatively small compared to the mOrange-tag and are expected to be imported into the mitochondrion, we anticipated that interference of the tag with import or proper folding could lead to the observed weak and undefined signals. To assess this, we selected two similarly small and abundantly expressed targets, PBANKA_0310100 (127 amino acids) and PBANKA_1024800 (144 amino acids), to test different tags. For PBANKA_0310100, which was already successfully localized to the mitochondrion, these included tags consisting of only the fluorescent protein mOrange or only the 3xHA epitope tag, either with or without a linker sequence. For PBANKA_1024800, we also included the combined mCherry-cMyc tag that has previously been used successfully to tag mitochondrial proteins (17). We found that the fluorescent protein tag indeed interfered with the localization of PBANKA_1024800, while the 3xHA tag, with or without a linker, supported a mitochondrial localization (Supplemental Fig 5B). Surprisingly, using the mOrange-tag without the subsequent HA-tag also led to an undefined rather punctuate mislocalization of PBANKA_0310100 (Supplemental Fig 5B). Based on these observations, we decided to tag the remaining 11 proteins with a linker-3xHA-tag (Fig 3B, 3C and Supplemental Fig 4B). Five of these were also convincingly localized to the mitochondria (Fig 3B), while an additional three proteins showed very weak staining patterns suggestive of a possible mitochondrial localization (Fig 3C). The signals of the remaining three tagged proteins were unable to be distinguished from the background and therefore no subcellular localization could be assigned (Supplemental Fig 4B).

**Fig 3.**
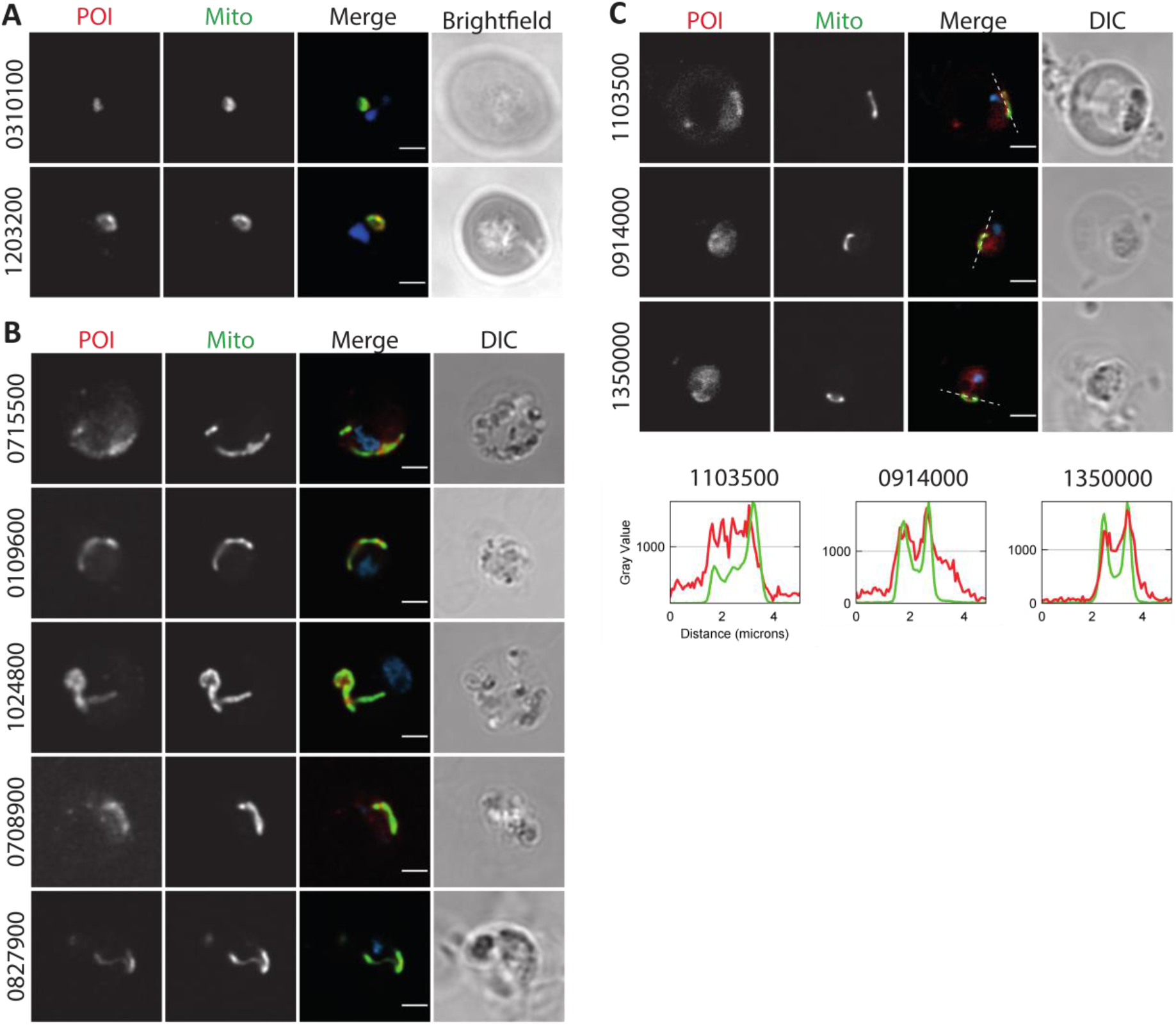
Validation of mitochondrial candidate proteins. Immunofluorescent analysis of tagged candidate proteins of interest (POI) in *P. berghei* parasites (PBANKA_ID indicated vertically). POI were stained with anti-HA antibody (red, first column), mitochondria were stained with anti-GFP antibody binding to the organelle marker (Mito, green, second column). The DNA was stained with DAPI (blue in merge). (A) Representative images of candidate proteins tagged with a mOrange-3xHA tag, imaged on a SP8 confocal microscope (Leica). (B-C) Representative images of candidate proteins tagged with a linker-3xHA tag, imaged on a LSM900 confocal microscope (Zeiss). (C) Images with the brightest signal for three lowly expressed candidate proteins. Pixel intensities of the POI and mitochondrial signal were normalized and plotted over the line indicated in the merge panel. Scale bar, 2 μm.

#### New mitochondrial proteins

A categorical representation of the identified proteins (Fig 2C) shows that 172 of the 445 proteins have a previous annotation of being mitochondrial as they are either part of the gold standard, have a mitochondrial GO term, or are described as putative mitochondrial protein. A smaller number of proteins, 30, were part of the negative set or have an apicoplast GO-term & no mitochondrial GO term. The remaining 243 proteins, of which 121 proteins are annotated as conserved hypothetical proteins, contain new mitochondrial candidates.

For 27 of the 121 conserved proteins with an unknown function, we found a mitochondrial ortholog using the sensitive homology detection tool HHpred (36) that can improve their annotation (Supplemental Table 2). For example, the human and yeast orthologs of PF3D7_0306000 (Bayesian rank 41) are both cytochrome b-c1 complex subunit 8, making it very likely that the *Plasmodium* protein is also cytochrome b-c1 complex subunit 8, as confirmed by recent complexome profiling (12). Other similar examples include PF3D7_1357600 (rank 285) that is likely a mitochondrial member of the 39S ribosomal protein family L53, and PF3D7_1309900 (rank 80) that is likely succinate dehydrogenase assembly factor 2.

Besides the possibility to improve annotations using well-characterized mitochondrial orthologs, the predicted 445 proteins contain biologically interesting proteins that can add new mitochondrial functions to *P. falciparum* and related species. Therewith, the prediction provides targets for experimental research to uncover new biology. These proteins can be divided into five categories: (i) proteins with known mitochondrial orthologs and an unknown mitochondrial role in *P. falciparum*, (ii) proteins without a mitochondrial ortholog but with homology to proteins with a known function and conservation of critical residues that suggest conservation of that function, (iii) proteins with homology to proteins with a known function from which residues known to be essential for catalytic activity or for binding substrates residues have been lost, (iv) new proteins of a known mitochondrial protein family and (v) proteins of a new mitochondrial protein family. Below, we describe several proteins per category, including their domain structure, the conservation of active site residues, their phylogenetic distribution and if available, the experimental confirmation of their mitochondrial localization

### Proteins with known mitochondrial orthologs and an unknown mitochondrial role in P. falciparum

PF3D7_0413500/PfPGM2 (rank 111) contains a PGM domain and appears to be orthologous to the human serine/threonine-protein phosphatase, PGAM5, of which the catalytic residue H105 (61) is conserved (62). PfPGM2 catalyzes the dephosphorylation of phosphorylated sugars and amino acids (62). In human cells, PGAM5 is localized to the mitochondrial outer membrane (63) where it plays a role in apoptosis, mitofission, and mitophagy (reviewed in (64)) among others via interactions with BCL2-like protein 1 (65). A previous study reported an unexpected cytoplasmic localization of this protein in *P. falciparum* and *P. berghei*, though in the absence of brightfield/DIC images or co-localization with marker proteins it is difficult to draw conclusions on the localization (62). Therefore, we tagged the *P. berghei* ortholog PBANKA_0715500 with a 3xHA tag and confirmed co-localization of PbPGM2 with our mitochondrial marker (Fig 3B). We also observed a weak non-mitochondrial background signal, which could indicate a potential dual localization. An important consideration in the interpretation of the previously reported localization of the GFP-tagged *P. berghei* protein is our observation that the use of a big tag such as GFP can interfere with the localization of mitochondrial proteins (Fig S5).

Originally, we flagged two additional proteins, PF3D7_0213200 (rank 169) and PF3D7_0611300 (rank 171), as interesting due to potential homology to BCL2, based on sequence similarity and a similar alpha-helical structure predicted with HHpred (36). PF3D7_0213200 was assigned an apicoplast GO-term, but in our analysis is a potential mitochondrial protein. Analysis with TMHMM (66) indicates that it probably is a transmembrane protein with an in-out topology. The protein PF3D7_0611300 already was assigned a mitochondrial GO-term based on a screening in *T. gondii* (67), but was, to our knowledge, not confirmed as mitochondrial localized with experiments characterizing this protein in *P. falciparum.* Although we rejected BCL2 homology for both proteins after a detailed examination of the residues conserved in the BCL2 family, we confirmed the mitochondrial localization of PF3D7_0213200 (PBANKA_0310100 in Fig 3A) and PF3D7_0611300 (PBANKA_0109600 in Fig 3B). Indeed, complexome profiling revealed that the latter is part of the ATP synthase complex (12).

### Proteins with a non-mitochondrial homolog and conservation of critical residues

PF3D7_0913400 (rank 30) is homologous to the bacterial elongation factor P. Although eukaryotes contain initiation factor 5A as a cytoplasmic elongation factor P homolog, this protein family was until now not observed in mitochondria. PF3D7_0913400 contains both the N-terminal KOW-like domain and the central P/YeiP domain of elongation factor P, but not the C-terminal OB domain. Elongation factor P plays a role in translation elongation and specifically in the translation of stretches of 2 or more prolines (68). A basic residue at position 34 of elongation factor P (numbering from the *Escherichia coli* PDB structure 3A5Z_F (69)) that after post-translational modification interacts with the CCA-end of the tRNA, is conserved in PF3D7_0913400. Notably, mitochondrial encoded *P. falciparum* cytochrome C oxidase subunit 1 contains a proline pair at position 135-136, making it a potential target for elongation factor P.

PF3D7_0812200 (rank 147) is homologous to the bacterial DegP protease family. It contains both a serine endoprotease domain and a PDZ domain. The DegP from *E. coli* functions in acid resistance in the periplasm. It is able to refold after acid stress and subsequently can cleave proteins misfolded due to acid stress (70). The active site residue (S210 in PDB structure 1KY9 (71)) is conserved in PF3D7_0812200, but residues involved in allosteric loop interactions, including R187 are not. Orthologs of PF3D7_0812200 appear, among the eukaryotes, limited to the Apicomplexa and Chromerida.

PF3D7_0503900 (rank 246) has a C-terminal dioxygenase domain (E= 1.3E-16). The closest homolog in human is the phytanoyl-CoA dioxygenase that resides in the peroxisome, an organelle absent from *Plasmodium*, where it is involved in α-oxidation of branched-chain fatty acids. Important residues in the active site that are involved in iron binding, H175, H177, H264 (72), are conserved in PF3D7_0503900, suggesting that it is a functional enzyme. Nevertheless, levels of sequence conservation are low (14% identity with the human phytanoyl-CoA dioxygenase) and we did not detect conservation of substrate binding sites with the phytanoyl-CoA dioxygenase. The co-localization analysis suggested a mitochondrial localization for PF3D7_0503900 (PBANKA_1103500 in Fig 3C).

PF3D7_1121500 (rank 307) contains a papain-like NLpC/P60 superfamily domain (Peptidase_C92 in PFAM). This protein family contains, among others, cysteine peptidases and amidases. Of the four residues essential for activity in BcPPNE from *Bacillus cereus* (73), the most similar experimentally characterized homolog of PF3D7_1121500, three are conserved (H49, E64, and Y164).

### Proteins with homology to proteins with a known function from which residues known to be essential for catalytic activity or for binding substrates residues have been lost

PF3D7_1004900 (rank 212) is homologous to the protein component of the signal recognition particle (SRP), a ribonucleoprotein that recognizes and targets specific proteins to the ER in eukaryotes. PF3D7_1004900 contains the C-terminal M domain (E= 5.8E-25), but not the other, SRP54 and SRP54_N domains of this protein. In SRP the M domain binds both the SRP RNA and the signal sequence of the target protein. Inspection of individual amino acids did not reveal conservation of RNA binding amino acids, e.g. the arginines in helix M4 of the M domain (74), are not conserved, but did reveal some conservation of hydrophobic amino acids lining the groove in which the signal peptide is located. These hydrophobic amino acids have been implicated in interactions with the signal peptide, specifically V323, I326, L329 in the M1 helix and L418 in the M5 helix (positions relative to *Sulfolobus solfataricus* structure 3KL4 (74, 75)), suggesting a possible conservation of interaction of PF3D7_1004900 with a hydrophobic α-helix. We experimentally confirmed the mitochondrial location of PF3D7_1004900 (PBANKA_1203200 in Fig 3A).

PF3D7_1246700 (rank 223) is a member of the pyridoxamine 5’-phosphate oxidase (PNPOx) like protein family that, besides pyridoxamine 5’-phosphate oxidase, also contains the general stress protein 26 family. The most similar experimentally characterized protein is general stress protein 26 from *Xanthomonas citri* encoded by gene *Xac2369*. The latter binds FAD and FMN but not pyridoxal 5’-phosphate, indicating that it does not function as a PNPOx (76). Relative to the unpublished structure of a protein from this family from *Nostoc punctiforme* (PDB: 2I02), that does contain an FMN, we did not observe conservation of the amino acids like W111, that line the FMN binding pocket, casting doubt of a function of PF3D7_1246700 that includes the binding of FMN.

### New members of mitochondrial protein families

PF3D7_1417900 (rank 109), PF3D7_1142800 (rank 172) and PF3D7_0927100 (rank 249) potentially contain a CHCH domain. They all have two pairs of cysteines that are very well conserved among their detectable homologs, and the nine amino acids between the cysteines are predicted to form an α-helix (36). Nevertheless, as nothing else but the cysteines appears to be conserved, the E-value with the PFAM CHCH domain entry, or with any known mitochondrial protein, is not significant (E>0.01). Jackhmmer analysis (77) detects homologs among the Apicomplexa, including a homolog in *Cryptosporidium muris*, but not outside this taxon. Known proteins with a CHCH domain are localized to the mitochondrial intermembrane space where they can participate in disulfide bond formation, suggesting that PF3D7_1417900, PF3D7_1142800, and PF3D7_0927100 might also be in the intermembrane space. Complexome profiling revealed that the first two are part of the ATP synthase complex (12). We tested PF3D7_1417900 and PF3D7_0927100 experimentally and confirmed them to be mitochondrial (PBANKA_1024800 and PBANKA_0827900 in Fig 3B respectively). Thus, based on our data and the complexome profiling data all three proteins are confirmed to be mitochondrial, increasing their likelihood that they indeed contain a CHCH domain.

*Plasmodium* contains a cytochrome C heme lyase (PF3D7_1224600, rank 129) and a cytochrome C1 heme lyase (PF3D7_1203600, rank 225), which are both essential during blood-stage development and have non-overlapping functions (78). We uncovered a third homolog in this family at rank 190, PF3D7_1121200 (E=0.0096, HHpred (36)). Although the homology covers domains II, III, and IV of heme lyases, of which domain II has been implicated in binding Heme (79), residues that have been implicated in heme binding, like H154 (coordinates for human protein (79)) and residues of domain I are not conserved in PF3D7_1121200. The protein also occurs in ciliates and dinoflagellates but has no orthologs outside of those taxa and therewith appears to be an alveolate-specific duplication.

### A new Apicomplexan mitochondrial protein family

We assessed to what extent *Plasmodium* mitochondrial proteins were part of families with multiple mitochondrial members that resulted from duplications (Supplemental Table 2). Such “intra compartmental protein duplications” have also increased the size of the human mitochondrial proteome (37). Among the mitochondrial homologs, we uncovered a new family in the Apicomplexa that in *Plasmodium* species has four representatives: PF3D7_0821900 (rank 209), PF3D7_1336100 (rank 258), PF3D7_1134400 (rank 280), and PF3D7_1228000 (rank 499). The family is characterized by a well-conserved WPP motif at its N-terminus (Supplemental Fig 7), and we therefore name it the WPP family. Although PF3D7_1228000 does not rank sufficiently high to fall within the 25% cFDR cut-off, all proteins of the family are likely mitochondrial given their overall high ranking and the tendency of members of protein families to be localized in the same cellular location. For the protein family itself, we did not detect homologs outside the Apicomplexa and *Vitrella*. In several species including *Plasmodium coatneyi* (PCOAH_00054040), *Babesia bovis* (BBOV_III010100), and *Theileria orientalis* (MACL_00002472), the ortholog of PF3D7_1228000 is predicted to be fused with an rRNA pseudourydilate synthase hinting at a potential function of this family in rRNA maturation. Nevertheless, there are no experimental data supporting those gene fusions. We unequivocally confirmed the mitochondrial location of PF3D7_0821900 (PBANKA_0708900 in Fig 3B) and the co-localization analysis also suggested a mitochondrial localization for PF3D7_1336100 and PF3D7_1134400 (PBANKA_1350000 and PBANKA_0914000 in Fig 3C respectively). PF3D7_1228000 was excluded from the analysis due to low transcript levels.

## Discussion

In this work, we ranked all nuclear encoded proteins of *P. falciparum* for their likelihood of being mitochondrially targeted and defined the top 445 proteins (cFDR 25%) as likely mitochondrial proteins (Supplemental Table 2). Despite the large variation between the datasets in the coverage of the nuclear encoded proteome and in the ability to identify mitochondrial proteins (Table 1, Fig 1), the probabilistic approach we used was able to integrate the data in an unbiased way by assessing the distribution of gold standard genes within each dataset during the scoring process.

### Quality of the predictions

Several observations indicate the high quality of the predictions. Firstly, the 10-fold cross-validation with an area under the curve of 0.959 shows the high predictive value and robustness of the prediction. Secondly, almost half of the predicted proteins have a previous annotation of being mitochondrial, even though most of these were not used in weighing the datasets. Thirdly, of thirteen proteins that we set out to validate experimentally, the seven for which we obtained a high enough expression to convincingly locate them in the cell, were all mitochondrial, while for three others the experimental data was consistent with a mitochondrial localization but equivocal due to low expression levels. Fourth, even though our top 445 contains nine proteins of the negative gold standard that was not manually supervised for all entries, manual examination of those nine showed that the majority are likely mitochondrial after all. Seven of them were detected in the hyperLOPIT study (22) and of those five were annotated as mitochondrial (legend Supplemental Table 2). Finally, several proteins that we predicted to be mitochondrial have, since we made those predictions, been shown to be mitochondrial using other methods. Specifically, complexome profiling of mitochondria-enriched fractions has elucidated the composition of mitochondrial complexes II, III, IV, and V (12). We identify in our top 445 proteins, six out of the seven members of complex II (four of which not in the GS or mitochondrial in Gene Ontology (GO)), ten out of the eleven members of complex III (none of which in the GS, three not in GO), seventeen out of the nineteen members of complex IV (none of which in the GS, thirteen not in GO), and twenty-one of the twenty three members of complex V (fifteen of which not in the GS, nine not in GO, Supplementary Tables 2 and 5). These results reinforce the robustness of the predictions and that the ranked list of the complete nuclear encoded *P. falciparum* proteome, which we name PlasmoMitoCarta, is a useful resource for *Plasmodium* researchers.

### New mitochondrial biology

The results of the integration provide ample material for exploring new mitochondrial biology as 185 out of the 445 proteins do not have a detectable mitochondrial ortholog in yeast, human, or plant. Some, however, do have homologs in bacteria that include conservation of critical residues, for example, the elongation factor P whose function, if confirmed, would be unique to mitochondria. They also include paralogs of the known mitochondrial proteins, like the twin Cx9C and heme lyase families, and a new mitochondrial protein family that contains a well-conserved WPP motif. Finally, they include PfPGM2, a protein that does have a human mitochondrial ortholog, but for which the experimental data thus far had however pointed to a cytoplasmic organization (62). The integrated ‘omics data and localization with a 3xHA tag do point at a mainly mitochondrial localization for a protein that in human is involved in apoptosis, mitofission, and mitophagy, processes about which little is known in *Plasmodium.* Such proteins provide leads to new mitochondrial biology.

As part of this study, we provide two other valuable resources. First, we included the ortholog information gathered for this study that can improve the annotation of *P. falciparum* proteins and provide researchers with additional hints for the function of a protein (Supplemental Table 2). In addition, the WICCA tool that we developed for this study is available online. It allows any researcher to assess co-expression of *P. falciparum* genes with a set of genes of four *Plasmodium* species across data from 83 microarray expression experiments, by weighing the datasets for their relevance to the set of genes. This can aid in the discovery of novel members of known pathways and molecular systems.

### Size of the mitochondrial proteome

We predict 445 mitochondrial proteins at the cut off, set at cFDR 25%. When using the posterior probabilities (2^mitochondrial score) of all *P. falciparum* genes to calculate the estimated number (E) of mitochondrial proteins by summing them all up (E=sum(P_posterior_/(P_posterior_+1), see ref (80) for a detailed explanation), we predicted the total size of the mitochondrial proteome to be 454 proteins. It therefore would appear that our prior odds of ~10%, which affects the total size calculation, overestimated the number of mitochondrial proteins. One can in principle lower the prior odds (see ref (80)) to 8.3% (412 proteins) such that the number is consistent with the estimated size of the proteome based on all data. However, the presence of five likely mitochondrial proteins in our gold negative sets leads to an underestimate of the size of the mitochondrial proteome. We decided not to manually weed out these inconsistencies, as this cannot be done systematically for all datasets, and applying this only to the apparent inconsistencies would result in circular arguments.

The predicted total size of 454 proteins is relatively small compared to the mitochondrial proteomes in human, plant, and yeast that consist of at least 900 proteins (5, 39, 40). However, small mitochondrial proteomes have been reported before, for example, 573 proteins in the distantly related ciliate *Tetrahymena thermophila* (81). In agreement with our findings, a recent study on the mitochondrial proteome of *T. gondii* that was not included in our integration reports a proteome of 421 proteins (16).

In conclusion, we combined the information of eight datasets with the curated gold standards to rank all nuclear encoded *P. falciparum* proteins on their likelihood of being mitochondrial in PlasmoMitoCarta. The value of this ranked proteome is shown by *in vivo* validation of top scoring proteins in the closely related species *P. berghei* and will provide an important resource for future investigations.

## Supporting information

Legends with the supplementray material

