## Supplementary figures and images for "A prioritized and validated resource of mitochondrial proteins in *Plasmodium* identifies leads to unique biology"

### Supplemental Figure 1

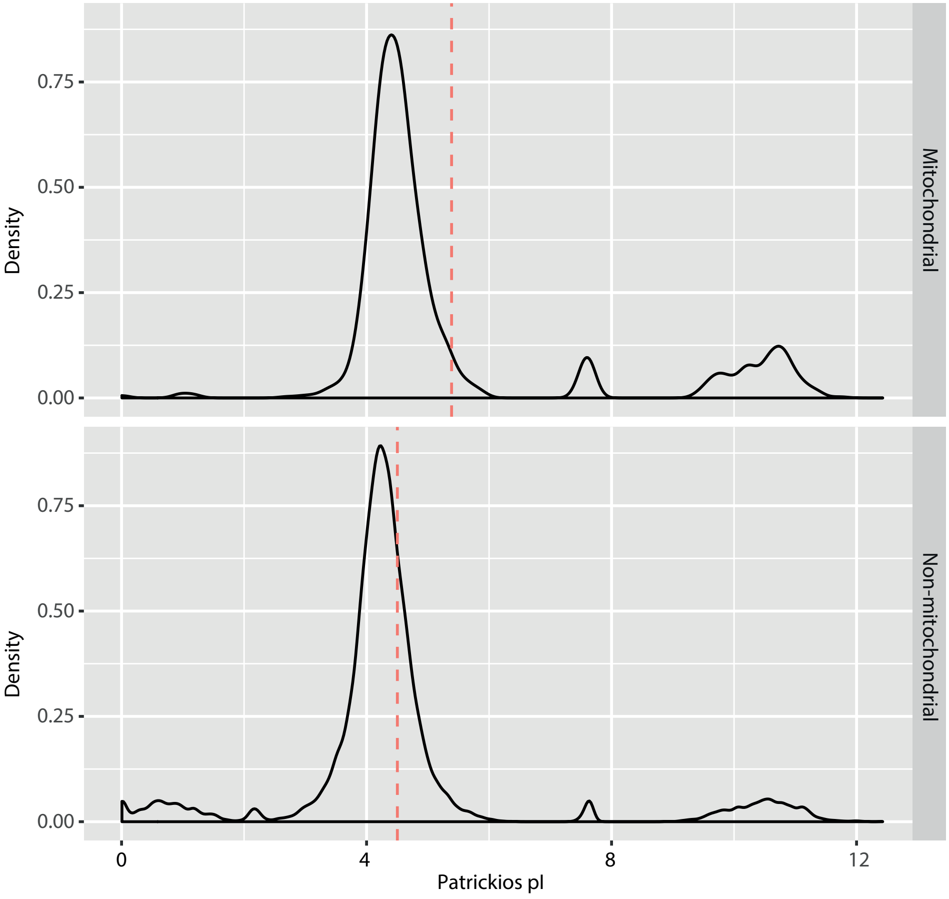

### Supplemental Figure 2

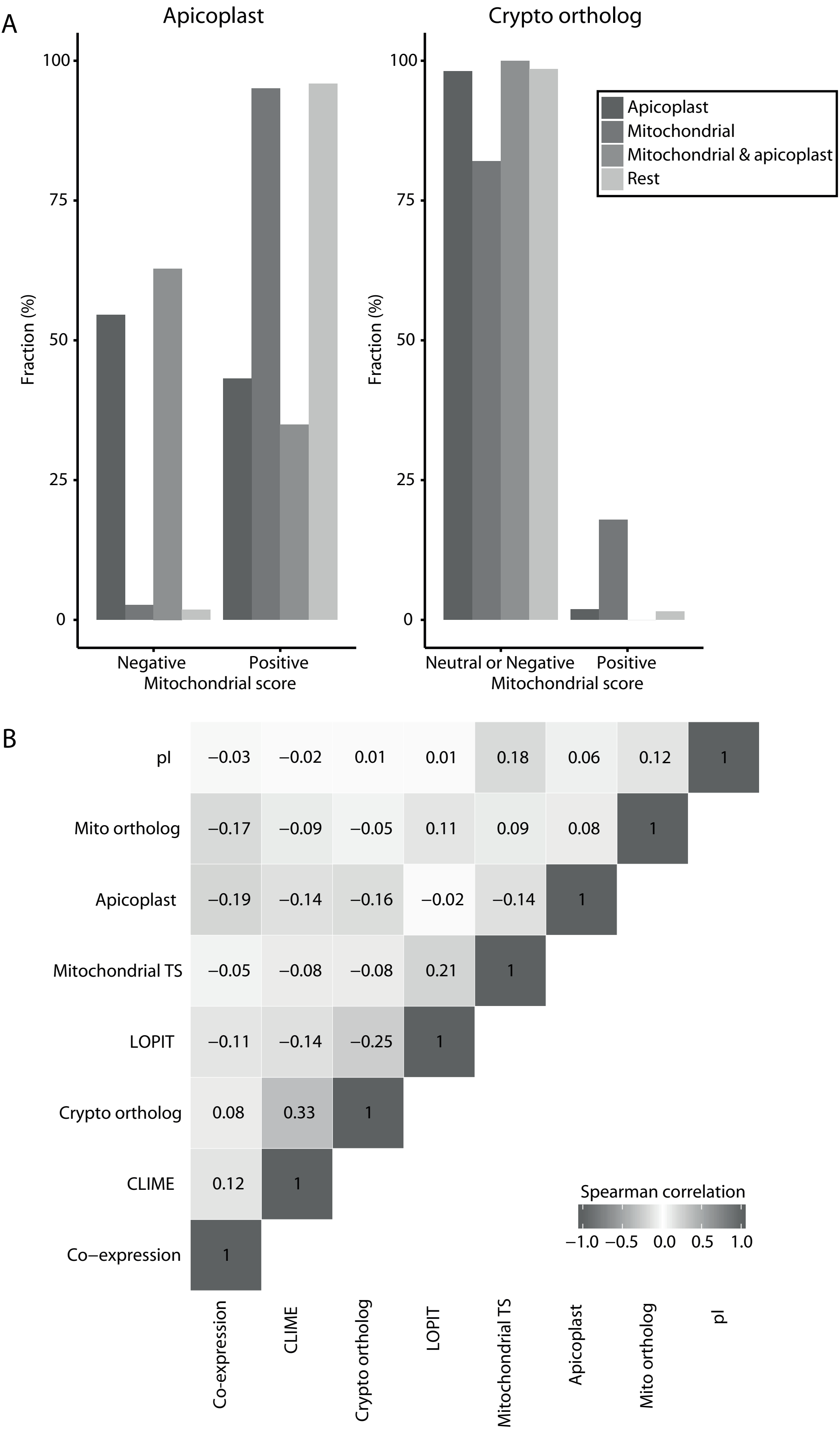

### Supplemental Figure 3

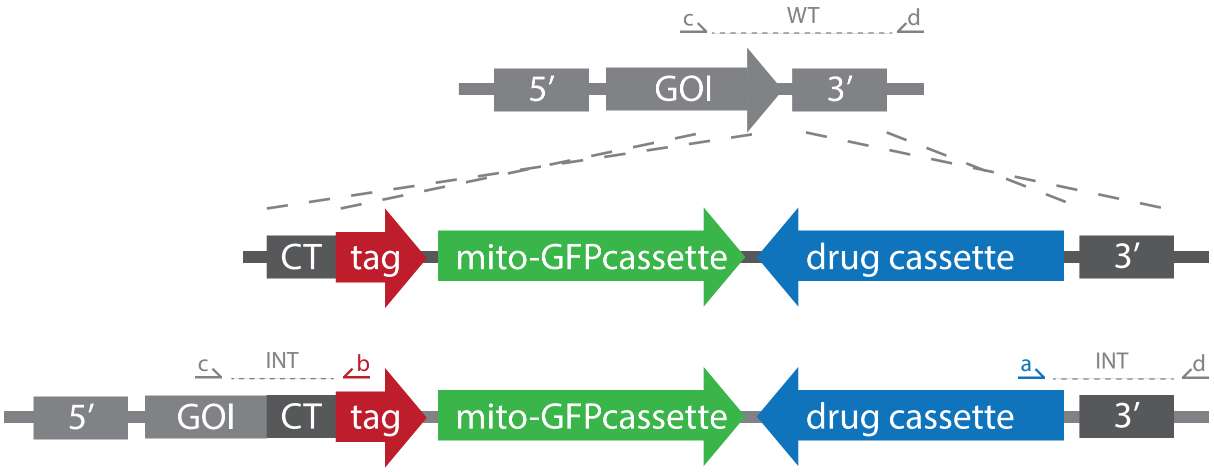

### Supplemental Figure 4

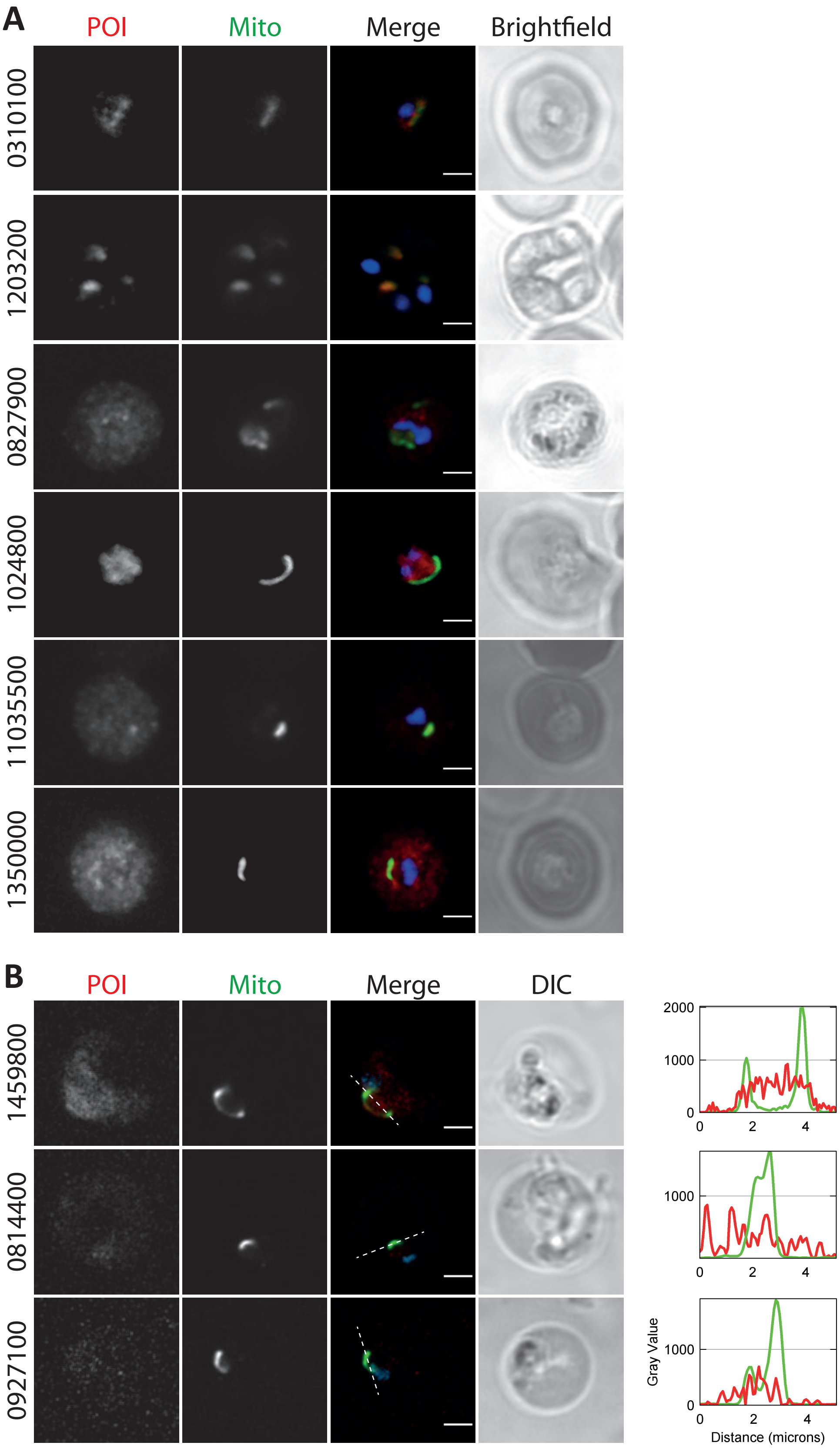

### Supplemental Figure 5

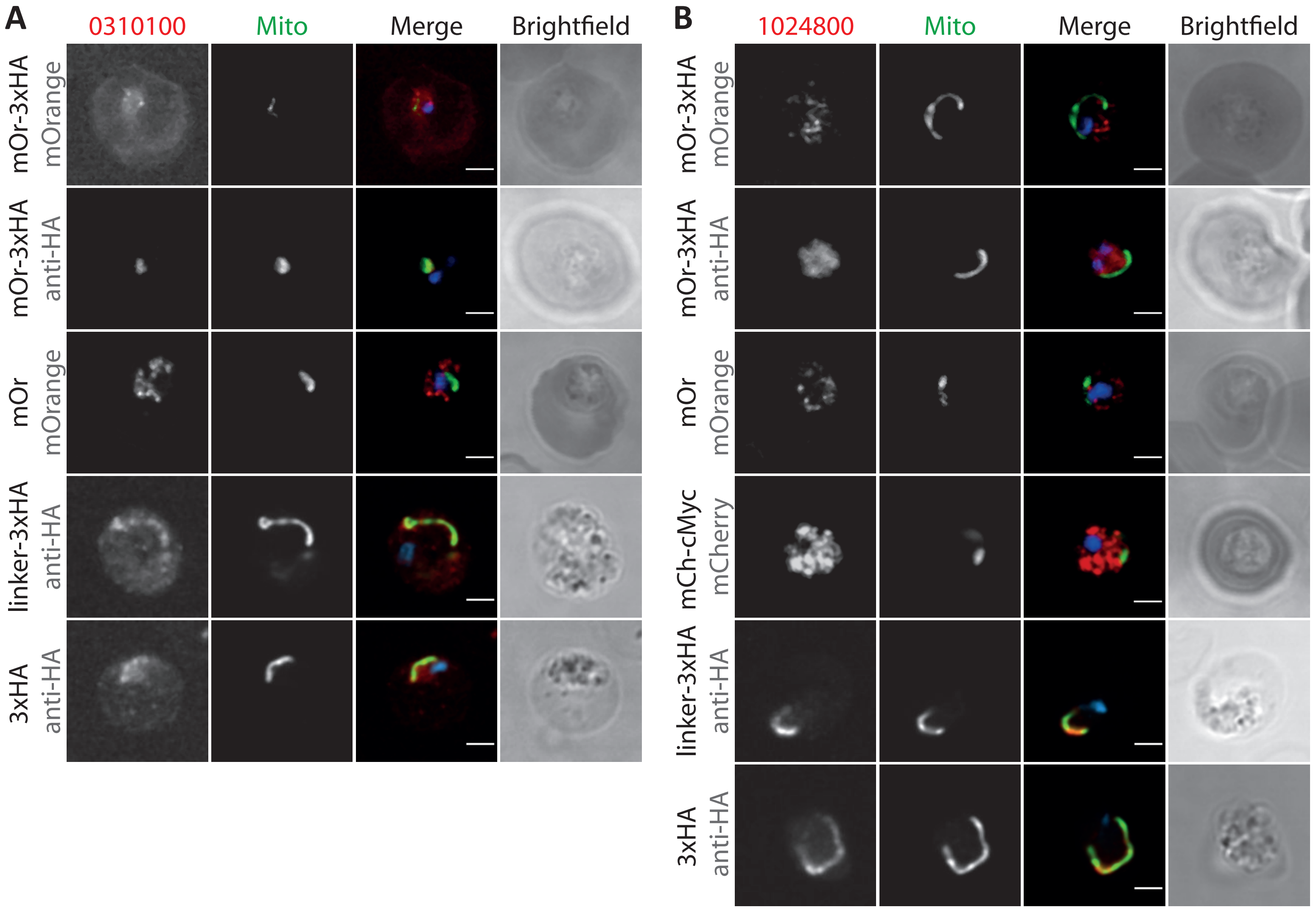

### Supplemental Figure 6

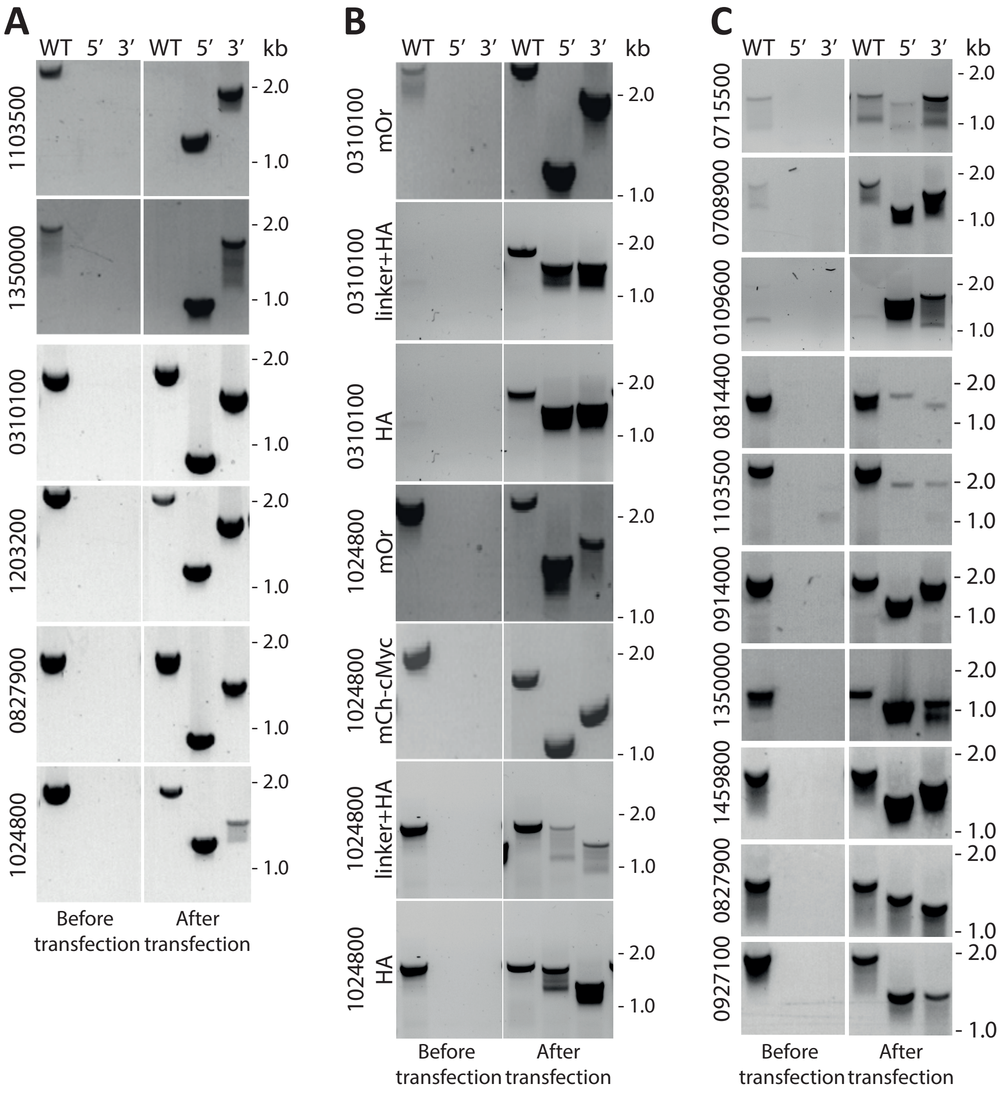

### Supplemental Figure 7

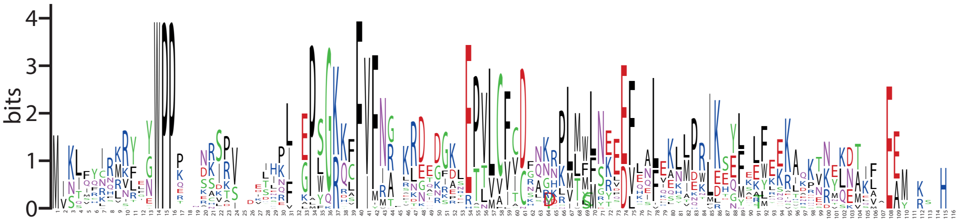
