## Supplementary material for "A prioritized and validated resource of mitochondrial proteins in *Plasmodium* identifies leads to unique biology": Legends with the supplementray material

**Supplement**

**Supplemental Table 1: Gold standards and alternative sets**

| Set | Subset | Gene IDs | | | | |
|---|---|---|---|---|---|---|
| Gold positive | Mitochondrial | PF3D7_0214200 | PF3D7_0607500 | PF3D7_0915300 | PF3D7_1134000 | PF3D7_1311000 |
| | | PF3D7_0217100 | PF3D7_0608310 | PF3D7_0925500 | PF3D7_1147700 | PF3D7_1311300 |
| | | PF3D7_0303700 | PF3D7_0709200 | PF3D7_1014700 | PF3D7_1215000 | PF3D7_1312400 |
| | | PF3D7_0415600 | PF3D7_0715500 | PF3D7_1015600 | PF3D7_1215300 | PF3D7_1312600 |
| | | PF3D7_0416600 | PF3D7_0727200 | PF3D7_1022500 | PF3D7_1217300 | PF3D7_1314600 |
| | | PF3D7_0420400 | PF3D7_0807400 | PF3D7_1028100 | PF3D7_1232200 | PF3D7_1320800 |
| | | PF3D7_0504600 | PF3D7_0808450 | PF3D7_1033000 | PF3D7_1233000 | PF3D7_1364900 |
| | | PF3D7_0518200 | PF3D7_0820700 | PF3D7_1034400 | PF3D7_1235700 | PF3D7_1365500 |
| | | PF3D7_0524700 | PF3D7_0823900 | PF3D7_1037300 | PF3D7_1239700 | PF3D7_1411600 |
| | | PF3D7_0603300 | PF3D7_0825200 | PF3D7_1117600 | PF3D7_1246100 | PF3D7_1428600 |
| | | PF3D7_0603700 | PF3D7_0915000 | PF3D7_1132900 | PF3D7_1310000 | |
| Gold negative | Apicoplast | PF3D7_0206100 | PF3D7_0409400 | PF3D7_0628800 | PF3D7_1124500 | PF3D7_1411400 |
| | | PF3D7_0208500 | PF3D7_0508800 | PF3D7_0711000 | PF3D7_1145500 | PF3D7_1440300 |
| | | PF3D7_0208600 | PF3D7_0522700 | PF3D7_0907900 | PF3D7_1209600 | PF3D7_1469000 |
| | | PF3D7_0302600 | PF3D7_0530200 | PF3D7_0908100 | PF3D7_1232100 | PF3D7_1469600 |
| | | PF3D7_0307400 | PF3D7_0602300 | PF3D7_0921400 | PF3D7_1251700 | |
| | | PF3D7_0312100 | PF3D7_0602400 | PF3D7_1005000 | PF3D7_1333200 | |
| | | PF3D7_0316900 | PF3D7_0604700 | PF3D7_1022800 | PF3D7_1357200 | |
| | | PF3D7_0408900 | PF3D7_0615100 | PF3D7_1103400 | PF3D7_1406600 | |
| | Non-mitochondrial | PF3D7_0729200 | PF3D7_0801900 | PF3D7_1123900 | PF3D7_1444300 | PF3D7_0802200 |
| | | PF3D7_1037100 | PF3D7_0802000 | PF3D7_1203500 | PF3D7_1445600 | PF3D7_0802800 |
| | | PF3D7_1205700 | PF3D7_0807900 | PF3D7_1212900 | PF3D7_1446500 | PF3D7_0902200 |
| | | PF3D7_1367700 | PF3D7_0810300 | PF3D7_1213800 | PF3D7_1447900 | PF3D7_0905100 |
| | | PF3D7_1037500 | PF3D7_0813000 | PF3D7_1219600 | PF3D7_1455100 | PF3D7_0936800 |
| | | PF3D7_0109800 | PF3D7_0914200 | PF3D7_1223400 | PF3D7_1461900 | PF3D7_1113700 |
| | | PF3D7_0110900 | PF3D7_0916100 | PF3D7_1249300 | PF3D7_1462000 | PF3D7_1116800 |
| | | PF3D7_0222100 | PF3D7_0925400 | PF3D7_1307800 | PF3D7_1468500 | PF3D7_1142400 |
| | | PF3D7_0302800 | PF3D7_0927600 | PF3D7_1317400 | PF3D7_0102200 | PF3D7_1203700 |
| | | PF3D7_0314400 | PF3D7_0931900 | PF3D7_1355700 | PF3D7_0202200 | PF3D7_1211600 |
| | | PF3D7_0315800 | PF3D7_1008900 | PF3D7_1401600 | PF3D7_0310400 | PF3D7_1221000 |
| | | PF3D7_0319000 | PF3D7_1009400 | PF3D7_1403600 | PF3D7_0416800 | PF3D7_1253200 |
| | | PF3D7_0319700 | PF3D7_1010300 | PF3D7_1412500 | PF3D7_0424400 | PF3D7_1333700 |
| | | PF3D7_0410300 | PF3D7_1013300 | PF3D7_1416500 | PF3D7_0501200 | PF3D7_1355300 |
| | | PF3D7_0606900 | PF3D7_1018200 | PF3D7_1423300 | PF3D7_0501300 | PF3D7_1418100 |
| | | PF3D7_0622800 | PF3D7_1033700 | PF3D7_1427200 | PF3D7_0507300 | PF3D7_1438900 |
| | | PF3D7_0711500 | PF3D7_1034900 | PF3D7_1441400 | PF3D7_0718100 | PF3D7_1457200 |
| | Non-apicoplast & non-mitochondrial Ralph Lab | PF3D7_0102500 | PF3D7_0423500 | PF3D7_0911900 | PF3D7_1129000 | PF3D7_1335300 |
| | | PF3D7_0109100 | PF3D7_0423700 | PF3D7_0917900 | PF3D7_1133400 | PF3D7_1335400 |
| | | PF3D7_0113700 | PF3D7_0423800 | PF3D7_0918000 | PF3D7_1141600 | PF3D7_1339900 |
| | | PF3D7_0202500 | PF3D7_0500800 | PF3D7_0924300 | PF3D7_1143100 | PF3D7_1342600 |
| | | PF3D7_0204100 | PF3D7_0501600 | PF3D7_0925700 | PF3D7_1202900 | PF3D7_1343100 |
| | | PF3D7_0204500 | PF3D7_0502400 | PF3D7_0927900 | PF3D7_1203000 | PF3D7_1346100 |
| | | PF3D7_0204700 | PF3D7_0503300 | PF3D7_0930300 | PF3D7_1216500 | PF3D7_1346700 |
| | | PF3D7_0206800 | PF3D7_0503600 | PF3D7_0932200 | PF3D7_1216600 | PF3D7_1347200 |

| | | | | | | |
|---|---|---|---|---|---|---|
| | | PF3D7_0206900 | PF3D7_0506200 | PF3D7_0932300 | PF3D7_1219000 | PF3D7_1355100 |
| | | PF3D7_0207000 | PF3D7_0507500 | PF3D7_0935800 | PF3D7_1222700 | PF3D7_1356900 |
| | | PF3D7_0207600 | PF3D7_0508000 | PF3D7_1014400 | PF3D7_1224300 | PF3D7_1364100 |
| | | PF3D7_0208900 | PF3D7_0511000 | PF3D7_1015900 | PF3D7_1227200 | PF3D7_1367500 |
| | | PF3D7_0214900 | PF3D7_0511200 | PF3D7_1016000 | PF3D7_1229400 | PF3D7_1369000 |
| | | PF3D7_0217500 | PF3D7_0524000 | PF3D7_1018300 | PF3D7_1232300 | PF3D7_1401400 |
| | | PF3D7_0220800 | PF3D7_0530900 | PF3D7_1027300 | PF3D7_1233600 | PF3D7_1403800 |
| | | PF3D7_0302100 | PF3D7_0604100 | PF3D7_1027700 | PF3D7_1234400 | PF3D7_1407800 |
| | | PF3D7_0303000 | PF3D7_0610400 | PF3D7_1028700 | PF3D7_1235200 | PF3D7_1408000 |
| | | PF3D7_0303400 | PF3D7_0616000 | PF3D7_1030200 | PF3D7_1240600 | PF3D7_1408100 |
| | | PF3D7_0304600 | PF3D7_0618500 | PF3D7_1031000 | PF3D7_1243700 | PF3D7_1409400 |
| | | PF3D7_0312400 | PF3D7_0628200 | PF3D7_1031200 | PF3D7_1246400 | PF3D7_1410400 |
| | | PF3D7_0315100 | PF3D7_0628300 | PF3D7_1035200 | PF3D7_1246900 | PF3D7_1436300 |
| | | PF3D7_0320400 | PF3D7_0702300 | PF3D7_1035500 | PF3D7_1247400 | PF3D7_1445400 |
| | | PF3D7_0320900 | PF3D7_0715000 | PF3D7_1036400 | PF3D7_1250100 | PF3D7_1448400 |
| | | PF3D7_0321400 | PF3D7_0717500 | PF3D7_1102800 | PF3D7_1252200 | PF3D7_1454400 |
| | | PF3D7_0324900 | PF3D7_0718300 | PF3D7_1104200 | PF3D7_1301700 | PF3D7_1455800 |
| | | PF3D7_0402300 | PF3D7_0722200 | PF3D7_1114100 | PF3D7_1302100 | PF3D7_1457000 |
| | | PF3D7_0406200 | PF3D7_0801300 | PF3D7_1115400 | PF3D7_1311800 | PF3D7_1457500 |
| | | PF3D7_0408600 | PF3D7_0814900 | PF3D7_1115700 | PF3D7_1314200 | PF3D7_1468400 |
| | | PF3D7_0408700 | PF3D7_0816400 | PF3D7_1116000 | PF3D7_1315100 | PF3D7_1471200 |
| | | PF3D7_0414900 | PF3D7_0823300 | PF3D7_1116100 | PF3D7_1315800 | PF3D7_1473700 |
| | | PF3D7_0415300 | PF3D7_0827900 | PF3D7_1116400 | PF3D7_1316200 | PF3D7_1475500 |
| | | PF3D7_0417100 | PF3D7_0830700 | PF3D7_1118400 | PF3D7_1318000 | PF3D7_1477700 |
| | | PF3D7_0418500 | PF3D7_0901000 | PF3D7_1121300 | PF3D7_1323700 | PF3D7_1479000 |
| | | PF3D7_0419700 | PF3D7_0903800 | PF3D7_1121700 | PF3D7_1332000 | |
| | | PF3D7_0423400 | PF3D7_0905400 | PF3D7_1124600 | PF3D7_1335100 | |
| Alternative positive | Mitochondrial GO-term | PF3D7_0106100 | PF3D7_0627400 | PF3D7_1125400 | PF3D7_1249500 | PF3D7_1430900 |
| | | PF3D7_0107400 | PF3D7_0708900 | PF3D7_1133300 | PF3D7_1249900 | PF3D7_1431000 |
| | | PF3D7_0108800 | PF3D7_0710900 | PF3D7_1137100 | PF3D7_1324400 | PF3D7_1431600 |
| | | PF3D7_0110200 | PF3D7_0716500 | PF3D7_1139700 | PF3D7_1325600 | PF3D7_1432100 |
| | | PF3D7_0202700 | PF3D7_0718400 | PF3D7_1141200 | PF3D7_1330600 | PF3D7_1434700 |
| | | PF3D7_0204800 | PF3D7_0815400 | PF3D7_1202200 | PF3D7_1336600 | PF3D7_1434800 |
| | | PF3D7_0204900 | PF3D7_0823700 | PF3D7_1202600 | PF3D7_1337000 | PF3D7_1435000 |
| | | PF3D7_0207200 | PF3D7_0825400 | PF3D7_1203600 | PF3D7_1340800 | PF3D7_1435800 |
| | | PF3D7_0207300 | PF3D7_0923600 | PF3D7_1208000 | PF3D7_1342100 | PF3D7_1439400 |
| | | PF3D7_0210100 | PF3D7_0923800 | PF3D7_1208300 | PF3D7_1342800 | PF3D7_1439600 |
| | | PF3D7_0212200 | PF3D7_0927800 | PF3D7_1208600 | PF3D7_1344100 | PF3D7_1441700 |
| | | PF3D7_0213700 | PF3D7_0928000 | PF3D7_1209800 | PF3D7_1345100 | PF3D7_1447300 |
| | | PF3D7_0302700 | PF3D7_0930900 | PF3D7_1209900 | PF3D7_1345200 | PF3D7_1453500 |
| | | PF3D7_0306400 | PF3D7_0933600 | PF3D7_1212000 | PF3D7_1345700 | PF3D7_1454500 |
| | | PF3D7_0311800 | PF3D7_1004800 | PF3D7_1212800 | PF3D7_1356200 | PF3D7_1454600 |
| | | PF3D7_0315500 | PF3D7_1008200 | PF3D7_1214200 | PF3D7_1368600 | PF3D7_1456100 |
| | | PF3D7_0416700 | PF3D7_1010000 | PF3D7_1214600 | PF3D7_1368700 | PF3D7_1458100 |
| | | PF3D7_0502200 | PF3D7_1012300 | PF3D7_1215200 | PF3D7_1404100 | PF3D7_1458700 |
| | | PF3D7_0513500 | PF3D7_1016400 | PF3D7_1223800 | PF3D7_1404900 | PF3D7_1459100 |
| | | PF3D7_0522500 | PF3D7_1021700 | PF3D7_1224600 | PF3D7_1408600 | PF3D7_1459300 |
| | | PF3D7_0523100 | PF3D7_1022900 | PF3D7_1227800 | PF3D7_1414900 | PF3D7_1460900 |

| | | | | | | |
|---|---|---|---|---|---|---|
| | | PF3D7_0531000 | PF3D7_1025600 | PF3D7_1230400 | PF3D7_1415800 | PF3D7_1462700 |
| | | PF3D7_0531200 | PF3D7_1103900 | PF3D7_1234900 | PF3D7_1416800 | PF3D7_1467700 |
| | | PF3D7_0603200 | PF3D7_1105800 | PF3D7_1235600 | PF3D7_1420300 | PF3D7_1470400 |
| | | PF3D7_0603500 | PF3D7_1108500 | PF3D7_1239100 | PF3D7_1424100 | PF3D7_1474100 |
| | | PF3D7_0611300 | PF3D7_1118200 | PF3D7_1239600 | PF3D7_1425200 | PF3D7_1475300 |
| | | PF3D7_0616800 | PF3D7_1123700 | PF3D7_1241600 | PF3D7_1426900 | |
| | | PF3D7_0617000 | PF3D7_1124700 | PF3D7_1242700 | PF3D7_1427600 | |
| | | PF3D7_0622600 | PF3D7_1125100 | PF3D7_1242900 | PF3D7_1428700 | |
| | | PF3D7_0623500 | PF3D7_1125300 | PF3D7_1245400 | PF3D7_1429700 | |
| Alternative negative | Apicoplast | PF3D7_0305000 | PF3D7_0607300 | PF3D7_1020800 | PF3D7_1234600 | PF3D7_1440200 |
| | | PF3D7_0314300 | PF3D7_0704900 | PF3D7_1021300 | PF3D7_1239500 | PF3D7_1440800 |
| | | PF3D7_0416100 | PF3D7_0716600 | PF3D7_1031400 | PF3D7_1318200 | PF3D7_1452300 |
| | | PF3D7_0503100 | PF3D7_0716900 | PF3D7_1034600 | PF3D7_1333000 | PF3D7_1467300 |
| | | PF3D7_0504000 | PF3D7_0815700 | PF3D7_1114800 | PF3D7_1345500 | |
| | | PF3D7_0504400 | PF3D7_0815900 | PF3D7_1223300 | PF3D7_1413500 | |
| | | PF3D7_0508300 | PF3D7_0816600 | PF3D7_1225100 | PF3D7_1430700 | |
| | | PF3D7_0529300 | PF3D7_0904700 | PF3D7_1232000 | PF3D7_1436800 | |
| | Non-mitochondrial | PF3D7_0815000 | PF3D7_0827100 | PF3D7_1138400 | PF3D7_0406400 | PF3D7_1023900 |
| | | PF3D7_1015200 | PF3D7_0912500 | PF3D7_1209000 | PF3D7_0410000 | PF3D7_1113100 |
| | | PF3D7_1126000 | PF3D7_0927700 | PF3D7_1222300 | PF3D7_0424500 | PF3D7_1121600 |
| | | PF3D7_1419800 | PF3D7_0931200 | PF3D7_1301400 | PF3D7_0501100 | PF3D7_1132800 |
| | | PF3D7_1420400 | PF3D7_1009600 | PF3D7_1309200 | PF3D7_0507200 | PF3D7_1138500 |
| | | PF3D7_0208800 | PF3D7_1012700 | PF3D7_1331600 | PF3D7_0516600 | PF3D7_1141800 |
| | | PF3D7_0304500 | PF3D7_1028400 | PF3D7_1332900 | PF3D7_0624600 | PF3D7_1149000 |
| | | PF3D7_0422300 | PF3D7_1030600 | PF3D7_1336900 | PF3D7_0731100 | PF3D7_1200800 |
| | | PF3D7_0511700 | PF3D7_1104000 | PF3D7_1350100 | PF3D7_0827800 | PF3D7_1206000 |
| | | PF3D7_0515900 | PF3D7_1105600 | PF3D7_1414400 | PF3D7_0902500 | PF3D7_1212100 |
| | | PF3D7_0520100 | PF3D7_1108200 | PF3D7_1417700 | PF3D7_0903700 | PF3D7_1246200 |
| | | PF3D7_0702400 | PF3D7_1108700 | PF3D7_1455000 | PF3D7_0904900 | PF3D7_1322100 |
| | | PF3D7_0708300 | PF3D7_1117300 | PF3D7_1466100 | PF3D7_0910000 | PF3D7_1355500 |
| | | PF3D7_0716300 | PF3D7_1124300 | PF3D7_1469200 | PF3D7_0919000 | PF3D7_1420700 |
| | | PF3D7_0810500 | PF3D7_1128200 | PF3D7_0104200 | PF3D7_0936300 | PF3D7_1463900 |
| | | PF3D7_0816900 | PF3D7_1132300 | PF3D7_0212100 | PF3D7_1001900 | PF3D7_1471100 |
| | | PF3D7_0824300 | PF3D7_1135100 | PF3D7_0309000 | PF3D7_1016300 | PF3D7_1478600 |
| | Non-apicoplast & non-mitochondrial Ralph Lab | PF3D7_0103800 | PF3D7_0506900 | PF3D7_0936400 | PF3D7_1211400 | PF3D7_1362400 |
| | | PF3D7_0106700 | PF3D7_0508500 | PF3D7_1001500 | PF3D7_1211900 | PF3D7_1366400 |
| | | PF3D7_0108300 | PF3D7_0512300 | PF3D7_1002200 | PF3D7_1212500 | PF3D7_1370300 |
| | | PF3D7_0112200 | PF3D7_0513800 | PF3D7_1003600 | PF3D7_1214100 | PF3D7_1401800 |
| | | PF3D7_0113900 | PF3D7_0515300 | PF3D7_1012200 | PF3D7_1218000 | PF3D7_1402300 |
| | | PF3D7_0114100 | PF3D7_0522600 | PF3D7_1012400 | PF3D7_1220900 | PF3D7_1404300 |
| | | PF3D7_0201900 | PF3D7_0523000 | PF3D7_1015800 | PF3D7_1228600 | PF3D7_1405600 |
| | | PF3D7_0202000 | PF3D7_0525800 | PF3D7_1016900 | PF3D7_1229100 | PF3D7_1406800 |
| | | PF3D7_0207500 | PF3D7_0525900 | PF3D7_1017100 | PF3D7_1229800 | PF3D7_1407000 |
| | | PF3D7_0207700 | PF3D7_0610800 | PF3D7_1023000 | PF3D7_1230700 | PF3D7_1407100 |
| | | PF3D7_0209000 | PF3D7_0612700 | PF3D7_1030900 | PF3D7_1243400 | PF3D7_1407900 |
| | | PF3D7_0209800 | PF3D7_0617800 | PF3D7_1032400 | PF3D7_1252100 | PF3D7_1409900 |
| | | PF3D7_0210600 | PF3D7_0617900 | PF3D7_1033200 | PF3D7_1301600 | PF3D7_1412000 |
| | | PF3D7_0210700 | PF3D7_0620400 | PF3D7_1035300 | PF3D7_1311900 | PF3D7_1417800 |

| | | | | |
|---|---|---|---|---|
| PF3D7_0211200 | PF3D7_0623000 | PF3D7_1035400 | PF3D7_1316600 | PF3D7_1423100 |
| PF3D7_0212600 | PF3D7_0707300 | PF3D7_1035600 | PF3D7_1317100 | PF3D7_1426200 |
| PF3D7_0214100 | PF3D7_0709000 | PF3D7_1035700 | PF3D7_1320000 | PF3D7_1430200 |
| PF3D7_0215800 | PF3D7_0717300 | PF3D7_1035800 | PF3D7_1320600 | PF3D7_1436600 |
| PF3D7_0302200 | PF3D7_0731500 | PF3D7_1036000 | PF3D7_1321900 | PF3D7_1439000 |
| PF3D7_0304000 | PF3D7_0808200 | PF3D7_1036300 | PF3D7_1323500 | PF3D7_1443000 |
| PF3D7_0308300 | PF3D7_0817700 | PF3D7_1038400 | PF3D7_1328800 | PF3D7_1444800 |
| PF3D7_0315200 | PF3D7_0817900 | PF3D7_1040200 | PF3D7_1331300 | PF3D7_1446200 |
| PF3D7_0320100 | PF3D7_0818900 | PF3D7_1105000 | PF3D7_1335900 | PF3D7_1446600 |
| PF3D7_0320800 | PF3D7_0822500 | PF3D7_1105100 | PF3D7_1337100 | PF3D7_1451600 |
| PF3D7_0323500 | PF3D7_0822600 | PF3D7_1108600 | PF3D7_1340900 | PF3D7_1452000 |
| PF3D7_0404700 | PF3D7_0824400 | PF3D7_1115800 | PF3D7_1341600 | PF3D7_1456800 |
| PF3D7_0404900 | PF3D7_0831800 | PF3D7_1116700 | PF3D7_1343000 | PF3D7_1459900 |
| PF3D7_0405200 | PF3D7_0902100 | PF3D7_1118600 | PF3D7_1345600 | PF3D7_1460100 |
| PF3D7_0405900 | PF3D7_0909300 | PF3D7_1129100 | PF3D7_1346800 | PF3D7_1460600 |
| PF3D7_0406100 | PF3D7_0910600 | PF3D7_1130800 | PF3D7_1347700 | PF3D7_1462800 |
| PF3D7_0414600 | PF3D7_0911300 | PF3D7_1136900 | PF3D7_1349300 | PF3D7_1465500 |
| PF3D7_0424100 | PF3D7_0920700 | PF3D7_1141900 | PF3D7_1353200 | PF3D7_1474900 |
| PF3D7_0424200 | PF3D7_0923300 | PF3D7_1144900 | PF3D7_1353600 | PF3D7_1477300 |
| PF3D7_0424300 | PF3D7_0929400 | PF3D7_1145400 | PF3D7_1361100 | |
| PF3D7_0505800 | PF3D7_0935900 | PF3D7_1147800 | PF3D7_1361900 | |

**Supplemental Fig 1. Human mitochondrial proteins on average have a slightly higher isoelectric point (pI) compared to the non-mitochondrial remainder of the human proteome.** Density plots over the Patrickios pI retrieved from a proteome isoelectric point database (1) of the human mitochondrial proteome (upper panel) (2) and the non-mitochondrial remainder of the proteome (lower panel). Red dotted lines indicate the mean.

**Supplemental Fig 2. Features of the input datasets.** (A) Barplots that visualize the fraction of apicoplast proteins, mitochondrial proteins, dual localized mitochondrial and apicoplast proteins, and the remainder of the proteome per bin for the datasets ATS and *Cryptosporidium* ortholog. In this figure localization categories were based on mitochondrial and apicoplast associated GO-terms and not on gold or alternative standards. (B) Heatmap depicts the pairwise Spearman correlation coefficients between the input datasets used to predict the mitochondrial proteome.

**Supplemental Table 2: Prediction results.**
See separate excel file called ‘Supplemental Table 2’. Note that *T. gondii* orthologs of PF3D7_0816400, PF3D7_0927900, PF3D7_0924300, PF3D7_1008900 and PF3D7_0409400, which are part of the negative gold standard, have been identified in LOPIT studies as being mitochondrial.

**WeIghted Co-expression Calculation tool for plAsmodium genes (WICCA).** WICCA was written in R (3) (v3.3.1). The raw supplemented materials of 83 microarray experiments (12

*P. berghei*, 2 *P. chabaudi*, 66 P*. falciparum and*, 3 *P. vivax*) were retrieved from the Gene Expression Omnibus (GEO) (4) on 14 December 2016. The GEO ID's of the datasets can be found in Supplemental Table 3. The raw data was further processed using the R package limma (5) to correct for background noise and to perform a quantile normalization to remove systematic bias. Technical replicates were averaged and probe identifiers were re-mapped to the up-to-date gene identifiers via the platform's own annotation libraries and the PlasmoDB website (6). As we want to use all 83 datasets for each of the four species mentioned above, the ortholog group information from PlasmoDB (6) was used to remap each dataset to the other three species. Pearson's correlation coefficients were pre-calculated for every gene combination in all datasets as a measure of co-expression. The mean of these coefficients per dataset was calculated and used as background co-expression signal.

**Supplemental Table 3: Gene expression datasets used in the WICCA tool.**

| Species | GEO ID |
|---|---|
| *Plasmodium berghei* | GSE5672 |
| *Plasmodium berghei* | GSE9497 |
| *Plasmodium berghei* | GSE16259 |
| *Plasmodium berghei* | GSE34806 |
| *Plasmodium berghei* | GSE34877 |
| *Plasmodium berghei* | GSE52859 |
| *Plasmodium berghei* | GSE53246 |
| *Plasmodium berghei* | GSE58580 |
| *Plasmodium berghei* | GSE64887 |
| *Plasmodium berghei* | GSE65032 |
| *Plasmodium berghei* | GSE80015 |
| *Plasmodium berghei* | GSE83667 |
| *Plasmodium chabaudi* | GSE33333 |
| *Plasmodium chabaudi* | GSE80015 |
| *Plasmodium falciparum* | GSE2878 |
| *Plasmodium falciparum* | GSE4582 |
| *Plasmodium falciparum* | GSE8099 |
| *Plasmodium falciparum* | GSE9152 |
| *Plasmodium falciparum* | GSE9724 |
| *Plasmodium falciparum* | GSE9853 |
| *Plasmodium falciparum* | GSE9868 |
| *Plasmodium falciparum* | GSE10022 |
| *Plasmodium falciparum* | GSE11763 |
| *Plasmodium falciparum* | GSE13578 |
| *Plasmodium falciparum* | GSE14524 |
| *Plasmodium falciparum* | GSE16259 |
| *Plasmodium falciparum* | GSE18075 |
| *Plasmodium falciparum* | GSE19010 |
| *Plasmodium falciparum* | GSE24416 |
| *Plasmodium falciparum* | GSE25642 |
| *Plasmodium falciparum* | GSE25878 |
| *Plasmodium falciparum* | GSE25879 |
| *Plasmodium falciparum* | GSE28625 |
| *Plasmodium falciparum* | GSE28701 |
| *Plasmodium falciparum* | GSE28990 |
| *Plasmodium falciparum* | GSE29874 |

| | |
|---|---|
| *Plasmodium falciparum* | GSE30869 |
| *Plasmodium falciparum* | GSE31829 |
| *Plasmodium falciparum* | GSE32211 |
| *Plasmodium falciparum* | GSE33605 |
| *Plasmodium falciparum* | GSE33764 |
| *Plasmodium falciparum* | GSE33795 |
| *Plasmodium falciparum* | GSE33796 |
| *Plasmodium falciparum* | GSE33797 |
| *Plasmodium falciparum* | GSE33826 |
| *Plasmodium falciparum* | GSE33834 |
| *Plasmodium falciparum* | GSE33835 |
| *Plasmodium falciparum* | GSE33836 |
| *Plasmodium falciparum* | GSE35732 |
| *Plasmodium falciparum* | GSE35949 |
| *Plasmodium falciparum* | GSE39238 (run on two platforms with different probes, so counted as two datasets) |
| *Plasmodium falciparum* | GSE39485 |
| *Plasmodium falciparum* | GSE41567 |
| *Plasmodium falciparum* | GSE44127 |
| *Plasmodium falciparum* | GSE47349 |
| *Plasmodium falciparum* | GSE47579 |
| *Plasmodium falciparum* | GSE47611 |
| *Plasmodium falciparum* | GSE52030 |
| *Plasmodium falciparum* | GSE53176 |
| *Plasmodium falciparum* | GSE54806 |
| *Plasmodium falciparum* | GSE56329 |
| *Plasmodium falciparum* | *GSE57748* |
| *Plasmodium falciparum* | GSE59015 |
| *Plasmodium falciparum* | GSE59097 |
| *Plasmodium falciparum* | GSE59098 |
| *Plasmodium falciparum* | GSE61536 |
| *Plasmodium falciparum* | GSE62364 |
| *Plasmodium falciparum* | GSE64887 |
| *Plasmodium falciparum* | GSE64688 |
| *Plasmodium falciparum* | GSE64690 |
| *Plasmodium falciparum* | GSE66669 (Split into three datasets of equal size due to high number of samples) |
| *Plasmodium falciparum* | GSE72578 |
| *Plasmodium falciparum* | GSE72579 |
| *Plasmodium falciparum* | GSE72695 |
| *Plasmodium falciparum* | GSE75295 (run on two platforms with different probes, so counted as two datasets) |
| *Plasmodium falciparum* | GSE83667 |
| *Plasmodium vivax* | GSE11075 |
| *Plasmodium vivax* | GSE12174 |
| *Plasmodium vivax* | GSE55644 |

The user can enter a list of query genes for WICCA to calculate their co-expression and find potentially other genes of interest that are co-expressed with the input genes. The Pearson correlation between each gene and each query gene was calculated for every dataset, background co-expression signal was subtracted. The mean of these values per gene in each dataset was taken to obtain one background-corrected value per gene for each dataset.

WICCA assumes that the query genes come from a common pathway or share a functional relationship and therefore the pair-wise correlation between these query genes influences the weight that each of the 83 datasets ultimately get. A higher weight should be given to datasets that show high correlation between the query genes and less weight to datasets that do not. Weighing was performed by calculating a weighing score ($\omega_d$) for each dataset with the following formula:

$$\omega_d = \frac{\sum_1^i \sum_1^j r_{x_{1..i},\ y_{1..j}}}{k^2 - k}$$

Where *r* is the Pearson's correlation coefficient of the expression values of gene *x* and *y*, *k* is the number of query genes, with *i* and *j* equal in size to *k* used here to distinguish between different combinations of query genes. The sum of the weights of all datasets is set to 1. The calculated weight for each dataset thus ranges between 0 and 1 and is a measure of how well the entered query genes co-express with each other within that particular dataset.

For each dataset the weighing score is multiplied by the calculated background-corrected Pearson correlation values. The sum of all weighted co-expression values associated with a gene across all datasets was calculated to obtain one final weighted gene-specific co-expression score. In the final results genes are ranked on this weighted value, with genes showing high co-expression with the query genes occurring at the top of the ranking.

**Evolutionary inference network setup.** In order to calculate the CLIME scores from the 138 species listed in the database, we used a multilayer perceptron (MLP) with two hidden layers, with 16 and 4 hidden nodes. For node activation, the softplus function was used. As the network output, two scores were calculated; a positive and negative score (for resembling the positive or the negative set, respectively). The final score was calculated as positive/(positive + negative) to represent a relative score.

Using the alternative positive/negative set as training data, the error was back-propagated through the network using the gradient descent method, adjusting the weights in between layers. 1280 iterations were performed. The resulting network was used to calculate scores for the testing data, which were later evaluated using 4-fold cross-validation. The area under the mean receiver-operator curve was 0.76$\pm$0.05, indicating that the CLIME scores are useful predictors for mitochondrial localization. This evaluation was performed independently of the Bayesian 10-fold cross-validation.

**Generation of tagging plasmids.** The initial vector including an hDHFR drug-selectable cassette, a mitochondrial GFP marker cassette, and a mOrange-3xHA tag with spacer sequence was obtained through triple ligation of elements from three different vectors. The plasmid backbone was derived from the pBAT vector (GenBank JX099571 (7)) digested with XhoI and EcoRI (2.1 kb), the drug-selectable and mitochondrial GFP marker cassettes were derived from the mitochondrial co-localization vector (8) digested with PvuII and XhoI (6.9 kb), and the linker-mOrange-3xHA tag was synthesized (GeneArt/Life Technologies) and digested with EcoRI and PmlI (1.3 kb). For testing of the effect of different tags on proteins localization, the mOrange-3xHA tag was replaced by the original mCherry-3xMyc tag, a mOrange-only tag, and two 3xHA tags, with and without linker sequence. Gene-specific 5' and 3' homology regions containing the carboxy-terminal (CT) ends and 3' untranslated regions (UTR) were amplified from *P. berghei* strain ANKA gDNA and cloned into the respective plasmids, such that the tag was fused in frame with the protein coding sequence (Supplementary Table 4, TV). All constructs were verified by commercial Sanger sequencing. Supplemental Fig 3 depicts a schematic overview of the tagging strategy.

**Generation of recombinant parasite lines.** The transfection plasmids targeting PBANKA_0715500 were linearized using AhdI and PvuI, all other plasmids were linearized using ApaLI and AhdI. Transfections were performed as described previously (9). When parasitemias reached 1%, typically at days 6-9 after transfection, blood of the infected mice was harvested to make samples for microscopy and isolate gDNA for genotyping. Correct integration of the transfected vector was assessed by diagnostic PCR (Supplementary Figure 6 and Supplementary Table 4).

**Supplemental Fig 3. Schematic overview of tagging strategy.** For all transfections the same double cross-over homologous recombination strategy was applied resulting in endogenously and stably tagging of the gene of interest (GOI). Every transfection vector contained a drug-selectable and a mitochondrial GFP marker cassette and was integrated into the genome via the C-terminus (CT) and 3'UTR (3'). As a result, the proteins of interest are fused in frame to the various tags applied in this study as indicated in the text allowing co-localization studies with the mitochondrial GFP marker. Primers used for the diagnostic PCR specific for wild-type (WT) or successful integration (INT) are indicated (a-d) and correspondent to those listed in Supplementary Table 4.

**Supplemental Fig 4. Localization of selected predicted mitochondrial proteins in *Plasmodium berghei*.** Tagged proteins of interest (POI) were stained with anti-HA antibodies (red) and corresponding mitochondrial GFP markers were stained with anti-GFP antibodies (green). DNA was stained with DAPI (blue). (A) Representative images of candidate proteins tagged with a mOrange-3xHA tag, imaged on a Leica SP8 confocal microscope. (B) Representative images of candidate proteins tagged with a linker-3xHA tag, imaged on a Zeiss LSM900 confocal microscope. The histogram depicts normalized pixel intensities of the POI and mitochondrial marker plotted over the line indicated in the merge panel. Scale bars, 2 µm.

**Supplemental Fig 5. Effect of different tags on protein localization.** (A) PBANKA_0310100 and (B) PBANKA_1024800. The tag integrated in the target locus is depicted vertically in black and the visualized signal of the protein of interest in gray. mOrange and mCherry signals (red) and the corresponding mitochondrial GFP markers (green) were visualized after fixation without the use of immunofluorescence. The 3xHA tags were stained with anti-HA antibodies (red) and corresponding GFP markers were stained with anti-GFP antibodies (green). DNA was stained with DAPI (blue). Scale bars, 2 µm. For completeness the image of PBANKA_0310100 (visible in Fig 3A) was repeated in this figure.

**Supplemental Fig 6. Diagnostic PCRs of *Plasmodium berghei* transfection experiments.** Integration PCR results on genomic DNA from parasites before (left) and after (right) transfection using the primer combinations shown in Supplemental Fig. 3 and Supplemental Table 4. Vertical numbers refer to the respective PBANKA_IDs. A) Assessment of integration of mOrange-3xHA tag. B) Integration of multiple tags for PBANKA_0310100 and PBANKA_1024800. mOr, mOrange; HA, 3xHA; mCh, mCherry. C) Assessment of integration of linker-3xHA tag.

**Supplemental Table 4: Primers used to create plasmids and recombinant parasite lines.**

**General primers**

| Primer name | Sequence | Use |
|---|---|---|
| **MF-20i** | GTAAAACGACGGCCAG | SQ |
| **MR-invitrogen** | CAGGAAACAGCTATGACC | SQ |
| **R-HA-tag** | CCGAAAAAGTTAAAATTAATTTAC | SQ |
| **(a) F-Pb-pDHFR** | ATGAAATACCGCTCCATTTTTCC | GT |
| **(b) R-mFruit** | GCCCTTGCTCACACCG | GT |
| **(b) R-HA-2** | CCATGGTAAATGGATGATTCCCCTCTTTGC | GT |

**PBANKA_0715500: phosphoglucomutase-2**

| Primer name | Sequence | Use | WT† | Int§ |
|---|---|---|---|---|
| **CT-0715500-F-EcoRI** | TTAAATGAATTCAGAAAAAATCGAAATGATGAAAATTCC | TV | 543 | |
| **CT-0715500-R-SpeI** | AAAATAACTAGTAAAATATGTAACACTATTGAAAGGAAGAT | TV | | |
| **3'-0715500-F-XhoI** | TTTTAACTCGAGTGCACATTAGGTCGTTACTACG | TV | 512 | |
| **3'-0715500-R-KpnI** | TTTTATGGTACCTTTTTTTCCATTTTGCAATGAAGC | TV | | |
| **(c) INT-0715500-F** | TAAGTTACAGGAAAAAAAAGAAGGG | GT | 1325 | 1192 |
| **(d) INT-0715500-R** | CAATTGGTCAAAGGGAAGAGC | GT | | 1288 |

**PBANKA_0109600: ATP-synthase-associated protein**

| Primer name | Sequence | Use | WT† | Int§ |
|---|---|---|---|---|
| **CT-0109600-F-EcoRI** | TTTAAGAATTCATGGTGGTAGATTTTTCATTTTGCC | TV | 744 | |
| **CT-0109600-R-SpeI** | AATTTAACTAGTTTTCGTTGCGGGTATTAGCA | TV | | |
| **3'-0109600-F-XhoI** | AAATTTCTCGAGTTAACAACATGAAAACAAACTCATGG | TV | 494 | |
| **3'-0109600-R-KpnI** | AATTAAGGTACCCAAATTTCACCATATCAACCGC | TV | | |
| **(c) INT-0109600-F** | GGTAAGCAGCATCCAAAATGG | GT | 1940 | 1448 |
| **(d) INT-0109600-R** | TTTGCCATGGGTTCATAGAGG | GT | | 1644 |

**PBANKA_1024800: ATP synthase-associated protein**

| Primer name | Sequence | Use | WT† | Int§ |
|---|---|---|---|---|
| **CT-1024800-F-EcoRI** | TTTATGAATTCATGAAATTAACAACCTGATGAAATGC | TV | 604 | |
| **CT-1024800-R-SpeI** | TAATTAACTAGTCCATTTATAAGCTACAAAATTTTCCTTAA | TV | | |
| **3'-1024800-F-XhoI** | AAATTTCTCGAGAGCATATGGAATAAAATGATTTCTGC | TV | 567 | |
| **3'-1024800-R-KpnI** | AATTATGGTACCCGCTACCAAATTAATCAACCG | TV | | |
| **(c) INT-1024800-F** | TAAGCAAGTGTAAAAATATAGAAAACG | GT | 1607 | 1566/1063/1062*/1534**/1062*** |
| **(d) INT-1024800-R** | GGTTATAAAAAGAGACAATGTTGAGG | GT | | 1226 |

**PBANKA_0310100: conserved protein, unknown function**

| Primer name | Sequence | Use | WT† | Int§ |
|---|---|---|---|---|
| **CT-0310100-F-EcoRI** | ATAATTGAATTCCGATTTAAATTTCCAGGGTTTTGG | TV | 791 | |
| **CT-0310100-R-SpeI** | TTTTATACTAGTTTTAATTAATTCAATATATTTTTCATATGATGCA | TV | | |
| **3'-0310100-F-XhoI** | AAAATTCTCGAGCATGTTCAGGTAAAAATGTCAATGTAC | TV | 541 | |
| **3'-0310100-R-KpnI** | TTTTATGGTACCTACTTGCATGTTCTTGAACTAGG | TV | | |
| **(c) INT-0310100-F** | AATTTTACCAGTCGAACAAGAACC | GT | 1713 | 1383/877/877*/1351** |
| **(d) INT-0310100-R** | ATAAATAACGAAATATAGAATCAAGTTTCC | GT | | 1411 |

**PBANKA_0708900: conserved protein, unknown function**

| Primer name | Sequence | Use | WT† | Int§ |
|---|---|---|---|---|
| **CT-0708900-F-EcoRI** | TTTAAAGAATTCGACGAAGATGGTAAAGAAGAACC | TV | 493 | |
| **CT-0708900-R-SpeI** | AAATTAACTAGTTTCATGAAAGGACTTTCTATATTCTTC | TV | | |
| **3'-0708900-F-XhoI** | TTAATTCTCGAGGCCATTATTTCAATCAGCAAGC | TV | 549 | |
| **3'-0708900-R-KpnI** | AATTAAGGTACCTTATGTTTAGACGAAGACCAAATCC | TV | | |
| **(c) INT-0708900-F** | TAATTTTAGCTATTTCCAAAGGTATAGG | GT | 1625 | 1111 |
| **(d) INT-0708900-R** | CCTCACACCTCCAACGATTAC | GT | | 1321 |

**PBANKA_1459800: conserved *Plasmodium* protein – unknown function**

| Primer name | Sequence | Use | WT† | Int§ |
|---|---|---|---|---|
| **CT-1459800-F-EcoRI** | TTTAAAGAATTCGAATCCGGAGAAATACCACTACC | TV | 522 | |
| **CT-1459800-R-SpeI** | TTTTAATACTAGTTACTTTAATCCATGATATTTTGTCATCAATC | TV | | |
| **3'-1459800-F-XhoI** | ATTTTACTCGAGATTTTGAATAGTTCTCATTTGAGGG | TV | 698 | |
| **3'-1459800-R-KpnI** | AAAATTGGTACCTTGCCAATGAAATAATATCAGTAATACC | TV | | |
| **INT-1459800-F** | CGATAAAATTGATAGAAGTTAACCCC | GT | 1627 | 1262 |
| **INT-1459800-R** | CATCCAACTGATCACATAGTGC | GT | | 1427 |

**PBANKA_0814400: conserved *Plasmodium* protein, unknown function**

| Primer name | Sequence | Use | WT† | Int§ |
|---|---|---|---|---|
| **CT-0814400-F-EcoRI** | TTTAAAGAATTCGGCAAATTAAACAAGGTAGATTGG | TV | 547 | |
| **CT-0814400-R-SpeI** | TTAAATACTAGTTTTTTTTCTTAAAGTTGCTAATAATGATG | TV | | |
| **3'-0814400-F-XhoI** | TTTTAACTCGAGTTTATTATTAATTTATATTTTTATTTTCAACTAGC | TV | 554 | |
| **3'-0814400-R-KpnI** | TTTAATGGTACCGCCAACTTTTAAAGCACAACG | TV | | |
| **(c) INT-0814400-F** | TGTATGTATATGCATGCTTGTATGG | GT | 1457 | 1264 |
| **(d) INT-0814400-R** | GCCTATATCCAATCCTATTATCAATGG | GT | | 1196 |

**PBANKA_1203200: conserved protein, unknown function**

| Primer name | Sequence | Use | WT† | Int§ |
|---|---|---|---|---|
| **CT-1203200-F-EcoRI** | ATAAAAGAATTCATGTAAAATGCTTAATGAAATATGAAGG | TV | 664 | |
| **CT-1203200-R-SpeI** | TATTAATACTAGTTTTTTTAATTCTAGGTTTGTTTCTAGAAAG | TV | | |
| **3'-1203200-F-XhoI** | AATTATCTCGAGATTTCATAGTTTGCTTAACTTCAAAG | TV | 478 | |
| **3'-1203200-R-KpnI** | ATTATAGGTACCATAATGAACCATGGAATATAATAAATGATTC | TV | | |
| **(c) INT-1203200-F** | AATACAATGTGAAATTTTGCTTTTTTTCC | GT | 1802 | 1511/1002 |
| **(d) INT-1203200-R** | ATAATATTATAAGCCCATTTAGACTTTTAG | GT | | 1435 |

**PBANKA_0827900: conserved protein, unknown function**

| Primer name | Sequence | Use | WT† | Int§ |
|---|---|---|---|---|
| **CT-0827900-F-EcoRI** | TTATATGAATTCTTTGGAGGGGAAAATGTCGAG | TV | 555 | |
| **CT-0827900-R-SpeI** | AATAATACTAGTCTGATTTCCCTTGACTATTTCAAG | TV | | |
| **3'-0827900-F-XhoI** | AAATTACTCGAGTGTTGAAAAAGTCCCACCAAGC | TV | 432 | |
| **3'-0827900-R-KpnI** | TATATTGGTACCCCAGATGAAATGTTTTCTCTCTCC | TV | | |
| **(c) INT-0827900-F** | CCATCAAAGCTCATATTTCAAAG | GT | 1467 | 1293/784 |
| **(d) INT-0827900-R** | TTAATGGTCAAGTACCGGATGC | GT | | 1154 |

**PBANKA_1103500: conserved *Plasmodium* protein, unknown function**

| Primer name | Sequence | Use | WT† | Int§ |
|---|---|---|---|---|
| **CT-1103500-F-EcoRI** | TAATTTGAATTCCCCCACTATTAATGGATAAAGATGC | TV | 776 | |
| **CT-1103500-R-SpeI** | AATTTAACTAGTTATATATCGGCCTAGTGTATTAAAGAA | TV | | |
| **3'-1103500-F-XhoI** | TAATATCTCGAGTATTTTCTTTCATTTCGAACAAGTGG | TV | 710 | |
| **3'-1103500-R-KpnI** | AAATTTGGTACCTTTTTGCAGAGTAGCTTTATTCG | TV | | |
| **(c) INT-1103500-F** | ACTTCAAAAGACATTGGGATCG | GT | 2192 | 1596/1087 |
| **(d) INT-1103500-R** | CGATTTATTTATTACCTTTAACCATTTTGC | GT | | 1570 |

**PBANKA_0927100: conserved protein, unknown function**

| Primer name | Sequence | Use | WT† | Int§ |
|---|---|---|---|---|
| **CT-0927100-F-EcoRI** | TTAAAAGAATTCTCAGCATTATTTCGGCAAGC | TV | 720 | |
| **CT-0927100-R-SpeI** | TTATAAACTAGTAAAATAGTAATATTTAAAAAATTTAAAAATATTTTTTATATG | TV | | |
| **3’-0927100-F-XhoI** | TATTTTCTCGAGTAATTTTTTTTCCATTGTTGGGTTTGC | TV | 556 | |
| **3’-0927100-R-KpnI** | TATATTGGTACCCCCACAGCATAGAGTTTTGC | TV | | |
| **(c) INT-0927100-F** | ATTTTTTAACGTGGATTTTTGTCCTG | GT | 1986 | 1360 |
| **(d) INT-0927100-R** | TATAAAGCAGTAAAGGAAATGTTCG | GT | | 1352 |

**PBANKA_0914000: conserved protein, unknown function**

| Primer name | Sequence | Use | WT† | Int§ |
|---|---|---|---|---|
| **CT-0914000-F-EcoRI** | AATAATGAATTCGAATAATGAAAAAATACCATCAGAAACG | TV | 527 | |
| **CT-0914000-R-SpeI** | TTATTTACTAGTTGTTTTCCTGCTTCTAAACACACC | TV | | |
| **3’-0914000-F-XhoI** | TTAAATCTCGAGTATTGATCCATCTTCATTACTTAAACCT | TV | 764 | |
| **3’-0914000-R-KpnI** | ATTAATGGTACCAGGACAAAAATGAAAACGACAGC | TV | | |
| **(c) INT-0914000-F** | GATACTCGTTTTGATAATGAAATTTCG | GT | 1698 | 1207 |
| **(d) INT-0914000-R** | CTCGAATTATGCAGCGTACC | GT | | 1575 |

**PBANKA_1350000: conserved protein, unknown function**

| Primer name | Sequence | Use | WT† | Int§ |
|---|---|---|---|---|
| **CT-1350000-F-EcoRI** | TTTAAAGAATTCCAAATAGAAAAGGGATAAATACTGAGG | TV | 591 | |
| **CT-1350000-R-SpeI** | AAATTAACTAGTAATTTTTTTCGCAAATTCTTCGAG | TV | | |
| **3’-1350000-F-XhoI** | TTAATTCTCGAGTTTTACGTACCGCCATTTAACG | TV | 667 | |
| **3’-1350000-R-KpnI** | AATTAAGGTACCTGATAAAATTCCATATAAAAGTACCATGC | TV | | |
| **(c) INT-1350000-F** | TGTAAACTATAAAGCCATTTGAATATCC | GT | 1501 | 1263/754 |
| **(d) INT-1350000-R** | TTATGTTAATTTTTTTTTCTCTCAACTCC | GT | | 1314 |

SQ: primers used for SeQuencing of the integrated locus or fluorescent cassette.
TV: primers used for creation of Tagging Vectors.
GT: primers used for GenoTyping after integration.
†: Expected PCR product size on wild-type *P. berghei* strain ANKA gDNA performed with forward (c) and reverse (d) primers.
§: Expected PCR product size on gDNA of successful transgenic parasites tagged with linker-3xHA / mOrange-3xHA / mOrange* / 3xHA** / mCherry-cMyc***.

**Supplemental Fig 7. Sequence logo of the mitochondrial WPP family.** An alignment was made using Jackhmmer (10) on the RefSeq database, starting with PF3D7_0821900 and iterating until convergence. The alignment was subsequently pasted in the sequence logo tool https://weblogo.berkeley.edu/ to obtain a graphical representation of the levels of conservation.

**Supplemental Table 5: Numbers of proteins retrieved by mass spectrometry of mitochondrial complexes (11) and mitochondrial-specific localization.** The numbers of proteins are relative to the predicted mitochondrial proteome. There are e.g. 4 mitochondrial complexes proteins that were predicted in the mitochondrial proteome for which there were no HYPERlopit data and that also had no previous annotation as being mitochondrial.

| | TOTAL | PREDICTED MITO | NOT IN POS GS | NOT IN GO | LOPIT: MISSING | LOPIT: NOT MITO IN FINAL PREDICTION | LOPIT: MISSING OR NOT MITO | NOT IN GS, GO, LOPIT |
|---|---|---|---|---|---|---|---|---|
| **COMPLEX II** | 7 | 6 | 4 | 4 | 1 | 0 | 1 | 0 |
| **COMPLEX III** | 11 | 10 | 10 | 3 | 0 | 1 | 1 | 1 |

| | | | | | | | | |
|---|---|---|---|---|---|---|---|---|
| **COMPLEX IV** | 19 | 17 | 17 | 13 | 1 | 1 | 2 | 1 |
| **COMPLEX V** | 23 | 21 | 15 | 9 | 2 | 0 | 2 | 2 |
| **TCA** | 9 | 8 | 5 | 1 | 0 | 0 | 0 | 0 |
| **SUM** | 69 | 62 | 51 | 30 | 4 | 2 | 6 | 4 |
